# The architecture of brain co-expression reveals the brain-wide basis of disease susceptibility

**DOI:** 10.1101/2020.03.05.965749

**Authors:** CL Hartl, G Ramaswami, WG Pembroke, S Muller, G Pintacuda, A Saha, P Parsana, A Battle, K Lage, DH Geschwind

## Abstract

Gene networks have proven their utility for elucidating transcriptome structure in the brain, yielding numerous biological insights. Most analyses have focused on expression relationships within a circumspect number of regions – how these relationships vary across a broad array of brain regions is largely unknown. By leveraging RNA-sequencing in 864 samples representing 12 brain regions in a cohort of 131 phenotypically normal individuals, we identify 12 brain-wide, 114 region-specific, and 50 cross-regional co-expression modules. We replicate the majority (81%) of modules in regional microarray datasets. Nearly 40% of expressed genes fall into brain-wide modules corresponding to major cell classes and conserved biological processes. Region-specific modules comprise 25% of expressed genes and correspond to region-specific cell types and processes, such as oxytocin signaling in the hypothalamus, or addiction pathways in the nucleus accumbens. We further leverage these modules to capture cell-type-specific lncRNA and gene isoforms, both of which contribute substantially to regional synaptic diversity. We identify enrichment of neuropsychiatric disease risk variants in brain wide and multi-regional modules, consistent with their broad impact on cell classes, and highlight specific roles in neuronal proliferation and activity-dependent processes. Finally, we examine the manner in which gene co-expression and gene regulatory networks reflect genetic risk, including the recently framed omnigenic model of disease architecture.

## Introduction

Human neuropsychiatric diseases are genetically complex, mostly adhering to a polygenic architecture^1^ consisting of thousands of risk-conferring variants and genes.^2,3^ In contrast to purely Mendelian disorders – where generalizable mechanistic insight can be obtained from the analysis of a single gene – the etiology of complex genetic disorders is organized around functional groups of genes or pathways.^4^ Be they members of a protein complex, components of a signaling cascade, or a collection of critical genes converging on a biological process, genes within these groups are expected to be co-regulated so as to be expressed at the appropriate levels to permit the group or pathway to function consistently.^5,6,7^ Recent work provides strong evidence that RNA co-expression and protein-protein interaction (PPI) networks provide a powerful organizing framework for understanding how such groups of genes are organized, with predictive power to prioritize disease-associated variation in psychiatric^8,9,10^ and other polygenic disorders.^11,12,13,14,15,16^ In polygenic disorders, where thousands of genes are involved, this network framework aids in characterizing relevant biological pathways by subdividing genes into smaller, tractable and coherent sets of modules for experimental analysis.^17,18^

Genetic studies of neuropsychiatric and neurodegenerative disorders have identified non-coding genetic variation as the largest contributor to disease liability.^19,20^ However, the hundreds of risk genes involved in these disorders are expressed across multiple brain regions and cell types, each of which has different functional consequences and likely different relevance to disease.^21^ These observations highlight the importance of systems biology in grouping transcriptional activity into coherent functional groups, enabling a more comprehensible understanding of disease.^22^ Gene co-expression networks further this understanding by linking together genes which co-vary across prevalent cell types and cell states within tissue.^23,24^

To inform our understanding of molecular mechanisms in human brain, and their potential relevance to disease, we create an unbiased atlas of co-expression networks across 12 human brain regions.^25^ We compare different network construction methods and demonstrate that the co-expression relationships defined in these networks are robustly identified using alternative network methods and orthogonal brain data sets. Combined with previous networks built from fetal brain across developmental time-points,^26^ these networks comprise a new resource for understanding the convergent pathways, time-points, and brain regions affected by disease-associated variation.

We use this resource to address several core biological questions. First, we show that co-expression in the brain is hierarchically organized into signatures that vary from brain-wide, to multi-region and to region-specific. We demonstrate that brain-wide and cross-regional networks correspond to signatures of prevalent cell types and biological processes. Second, we show that region-specific modules capture regionally upregulated genes, and reflect more specialized cellular subtypes. We provide evidence that complex neuropsychiatric diseases such as autism spectrum disorder (ASD) and schizophrenia (SCZ) are influenced by a combination of both widespread and focal disruption to gene expression. For both ASD and SCZ, three major types of genetic/functional genomic signals: differential expression, rare high-impact variants, and common low-effect variants, converge on cross-regional networks that implicate neuronal and neural progenitor cell types. We also identify signals of region-specific disruption in these disorders, implicating additional cortex-specific components. Finally, we incorporate our gene networks into a model of genetic architecture, asking whether these co-expression and co-regulatory networks exhibit a core-periphery structure that follows the recently framed omnigenic hypothesis.^27^

## Results

### Building robust human co-expression networks

To explore the molecular anatomy of the human brain starting at the tissue level, we utilize RNA-sequencing data from the Genotype-Tissue Expression Consortium (GTEx), focusing on the 12 major brain regions profiled: Cerebellum (CBL), cerebellar hemisphere (CBH), dorso-lateral pre-frontal cortex (PFC), Brodman area 9 (BA9), Brodman area 24 (BA24), hippocampus (HIP), amygdala (AMY), hypothalamus (HYP), substantia nigra (SNA), nucleus accumbens (ACC), caudate nucleus (CDT), and putamen (PUT) (**figure 1a**).

**Figure 1:**
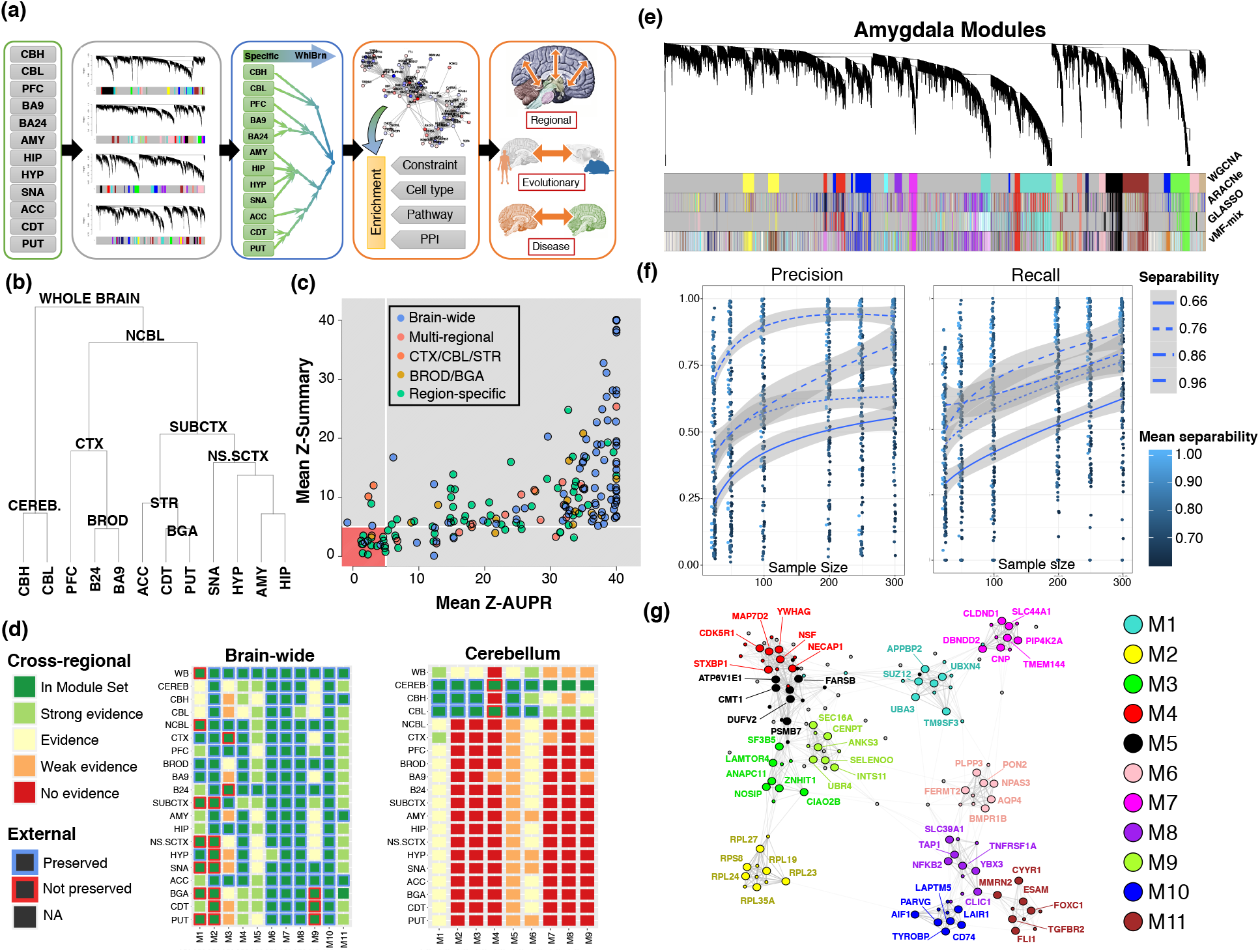
Human whole-brain co-expression atlas. (a) Overview of module construction (b) Hierarchical merging hierarchy based on median within-region expression (c) Module preservation in external datasets, with the not preserved (z < 5) region highlighted in red (d) Module evidence across all regions of the human brain, for brain-wide and cerebellar module sets. Strong evidence: z > 8, Evidence: z > 5, Weak evidence z > 3, No evidence z < 3. (e) Dendrogram from rWGCNA in amygdala, showing high degree of overlap between four methods of network construction and module identification. (f) Precision and recall of co-clustering a gene with the hub gene of its true module, as a function of module separability and sample size (g) Example hub gene network of whole-brain modules.

Technical artifacts induced by RNA quality, library preparation, and sequencing run are known to generate spurious gene-gene correlations while confounding true co-expression relationships.^28^ Taking advantage of the careful sample annotations and experimental protocols conducted in GTEx,^29^ we rigorously corrected for known technical covariates (**Methods**) to avoid these potential confounding factors. We evaluated the impact of using latent factors to correct for hidden confounders,^30^ and by examining the effect of latent variable correction on cell-type genes and pathway analysis, determined that it was removing significant biological signal, particularly regarding cell type (**figure S1; Suppl. Methods)**. This observation is consistent with recent results that caution against the use of latent factor correction when applying co-expression analysis in highly heterogeneous tissues such as brain.^31^

To ensure modules are not driven by brain-involved diseases or atypical sample outliers, we exclude individuals on the basis of their known medical conditions at time of death, principal-component outliers, and sample-sample connectivity outliers (**Methods**, **Suppl. Table 1**).^32^ In addition, we find that several outlier samples were taken from the same individuals, all of whom had a severe infection (sepsis, influenza, hepatitis, HIV), which is known to impact expression. We apply feature-selection and support vector machines to classify and remove these and any other samples where gene expression could be confounded by serious infectious disease (**Suppl. Methods**).

To reduce the impact of sample outliers, we use a bootstrapped-resampling version of weighted gene co-expression network analysis WGCNA (robust WGCNA, **Methods**)^33^ separately within each profiled region. To structure our analysis, we constructed a tissue hierarchy using median genome-wide expression, which produces a tissue grouping that reflects neuroanatomical and developmental regions (**figure 1b**). Using this hierarchy, we form a tree of consensus co-expression networks for each split, thereby generating co-expression modules for 20 hierarchical expression categories: 12 brain region specific categories (corresponding to each sampled region), 7 multi-regional categories (corresponding to multiple, structurally-linked regions, **figure 1b**), and a brain-wide category. The majority of the resulting modules are highly overlapping, therefore we group these modules hierarchically into groups of highly similar modules which we term “module sets” (**Methods**). In total, we identify 311 modules at all levels, of which 173/199 (87%) of the tissue-level modules are replicated with strong support in at least one other independent dataset (**figure 1c; Methods**). Finally, by using network preservation statistics on all samples within regions and meta regions, we verify that module sets are supported by strong evidence within their own regions and little evidence outside of them (**figure 1d**).

To test whether co-expression modules vary substantially by the method used for co-expression network construction, we build modules at the tissue level using three alternative approaches: ARACNe,^34^ PAM-guided graphical LASSO,^35^ and Fisher-von-Mises mixture modeling.^36^ We find that all methods show high pairwise clustering coefficients (**figure S1e**), and differ predominantly by module splitting (**figure 1e**). For instance, most of the differences between ARACNe clusters and WGCNA clusters come from large ARACNe modules represented as several separate WGCNA modules, consistent with prior comparisons that demonstrate this is due to WGCNA’s signed adjacency matrix carrying directional information, compared with ARACNe’s use of unsigned mutual information.^37^ In all cases, the vast majority of co-expression relationships identified by WGCNA are also co-clustered by each other method, suggesting that the results using WGCNA are generally robust to alternative methods of network construction (**figure S1e-h**).

We assessed the validity of our consensus-building approach by constructing whole-brain consensus networks using tensor decomposition,^38^ t-SNE,^39^ and the clustering algorithm dbSCAN,^40^ finding that all methods are in high correspondence (**Suppl. Methods, figure S1i-m**). We further apply a down-sampling approach to estimate overall power for module and hub detection (**Methods**), demonstrating power to comprehensively identify all module hub genes (**Figure S1n,o**). Module co-members are identified with expected lower, but reasonable precision (60%) recall (45%) and accuracy (55%) (**figure 1f,g**). Because thresholds for gene to module assignments can be arbitrary, we provide a table that lists genome-wide co-expression relationships for every gene-module pair (module eigengene correlation, kME; **Table S1**).

### Identifying brain-wide, regional, and tissue-specific modules

With the goal of establishing the regional specificity transcriptional programs throughout the brain, we utilize a hierarchical merging strategy using a combination of Jaccard similarity and module kME correlation to identify highly similar modules (**Methods**), thereby grouping modules into 48 *module sets*, 11 representing relationships present across the entire brain (Brain-Wide, or BW, **figure 1g**), and 38 consisting of those from regions within an established brain structure. We choose a simple naming convention by which a specific module is specified by its original tissue and module set: for instance, the hypothalamic module which is grouped into the whole-brain module set BWM10 is depicted as HYP-BWM10. As the organization of the regional transcriptome is necessarily complicated, we summarize our analyses at the whole-brain, multi-regional, and region-specific levels. To verify this approach in establishing shared and specific modules, we utilized the neighbor-based AUPR^41^ (**Methods**) to provide a degree of evidence for the top-level module set from the raw expression data (**figure 1d**), finding that brain-wide modules show evidence across all tissues, while the majority of region-specific modules show evidence only within that region (32/63) or adjacent regions (50/63). We find that the most distinct physiological regions, HYP, CBL, and SNA show the largest number of tissue-specific modules (**table S1**).

One module stands out in terms of preservation: Non-cerebellum (NCBL) M1 is only preserved in part of the striatum (caudate and nucleus accumbens) and telencephalon (amygdala and hippocampus), but not in putamen, cortex, or cerebellum; yet the module is preserved in several external microarray datasets. As the module eigengene is up-regulated in tissues adjacent to the ventricle and choroid plexus (CP - **figure 2a**), we hypothesized that this module could represent ependymal and choroid epithelial cells that are adjacent to the profiled regions. Indeed, we find that CP marker genes are enriched within the module (**figure 2a**), and, importantly, that regions lacking co-expression of NCBL.M1 module show low expression of CP cell markers. This finding demonstrates that our hierarchical approach identified the biologically correct placement of this cell type module, and the specificity of this module set corresponds directly to the presence or absence of the corresponding cell population.

**Figure 2:**
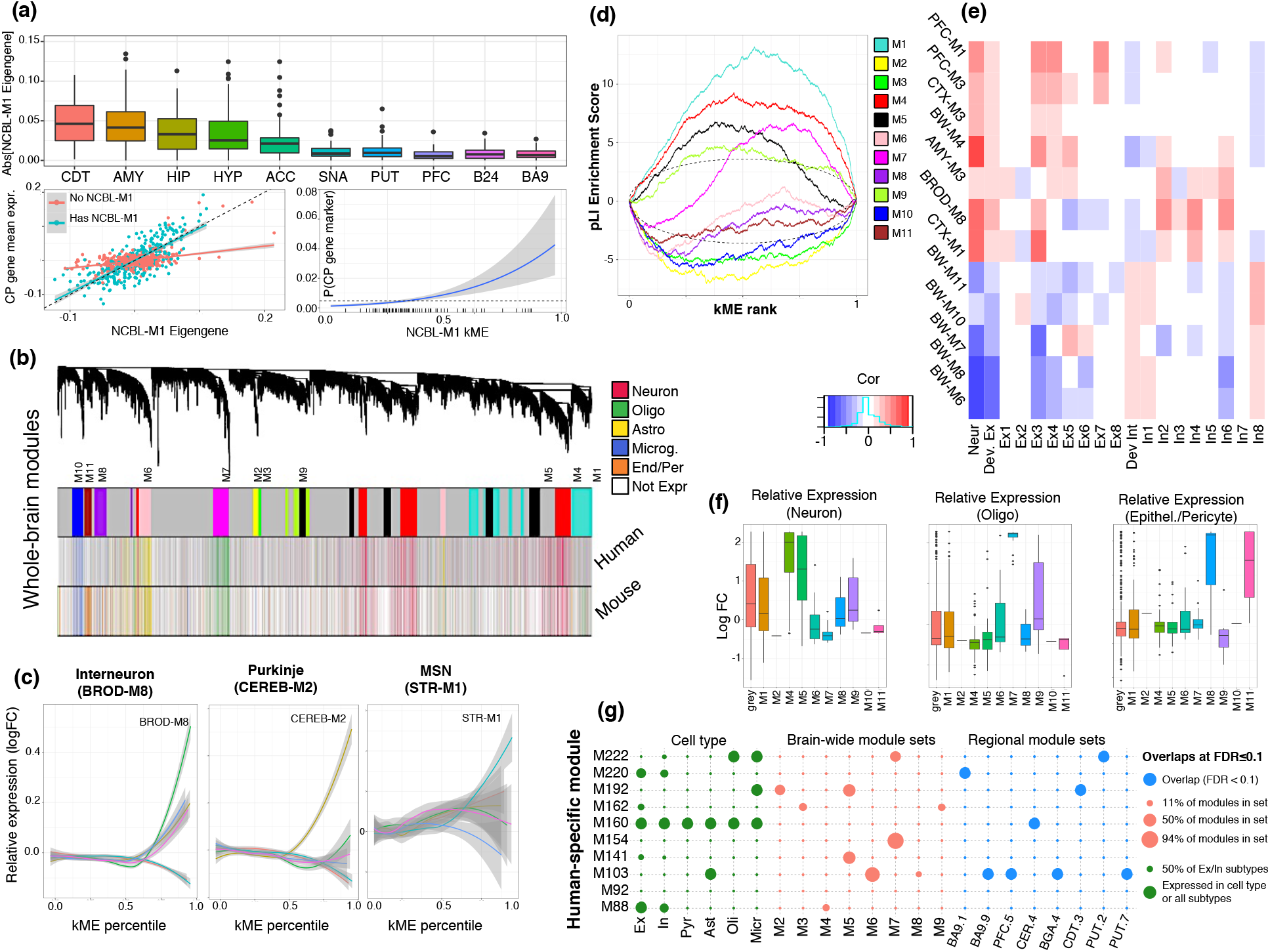
Cell-type heterogeneity relates to co-expression modules, gene intolerance, and evolution. (a) top: Absolute value of the eigengene of module NCBL.M1 plotted across regions, showing higher variance in regions adjacent to or accessible through ventricles. left: Expression of NCBL.M1 eigengene and mean expression of choroid-plexus marker genes in regions with and without an NCBL-M1 module. Right: Marginal probability of a gene being a choroid plexus marker, as a function of NCBL-M1 soft membership. (b) Brain-wide modules largely correspond to cell class. Top WGCNA dendrogram at the whole-brain level, colored by module, middle genes colored by cell type for which they are a marker in human and (bottom) mouse. (c) Relative expression of neuronal marker genes for modules BW-M4, BROD-M8, CEREB-M2, and STR-M1 within interneurons from cortical SC-sequencing, Purkinje neurons from cerebellar SC-seq, and medium spiny neurons from mouse striatal SC-seq, as a function of module kME. (d) GSEA enrichment plots for LoF-intolerant genes (pLI > 0.9) for all whole-brain modules. (e) Factorization-based decomposition of bulk expression. Correlations for BW, CTX, and PFC modules come from decomposition of DLPFC expression; AMY from decomposition of AMY bulk expression, and BROD from decomposition of B24 bulk expression. (f) lncRNA relative expression in single-cell data, grouped by the imputed module in riboZero RNAseq data from BA9. (g) Cell type expression and significant module overlaps for human differentially-expressed modules from Sousa et al. (17), with cell type assignments are as per (17).

### Module sets reflect common features and heterogeneity of brain cell types and processes

Cell type composition is a major driver of gene expression variance in tissue.^42,43^ We expect whole-brain co-expression modules to represent major cell classes (neurons, astrocytes, microglia, oligodendrocytes), and multi-regional or regional modules to represent more specialized cell subtypes. Using markers for primary brain cell types^44,45^ and cell subtypes^46^, we find that that modules at all levels significantly and specifically enrich for genes that correspond to a particular cell type. In particular, five of 11 whole-brain modules (M4, M6, M7, M10, M11) represent the 5 major brain cell classes (**figure 2b**), and two additional modules (M1, M8) capture cellular activity – neuronal differentiation and reactive gliosis, respectively (**figure S2a**). The region-specific modules BROD-M8, CEREB-M1, and STR-M2 correspond to interneurons, Purkinje cells, and medium spiny neurons, respectively; cell types that are either significantly enriched, or found only in these regions (**figure 2c**).

Genes intolerant to loss-of-function mutations in humans are known to be expressed disproportionately in brain,^47^ and particularly in cerebral cortex-expressed genes across all developmental periods,^48^ therefore we sought to link these genes to cell types and regions of the adult brain. We assessed whether any of our modules enrich for loss-of-function intolerant genes (defined as pLI^49^ > 0.9) and found that the whole brain module BW-M1 is the most significantly enriched for LoF intolerant genes, followed by module BW-M4 and another related module BW-M5 (**figure 2d, S2b**), all of which are neuronal. Notably only a single cluster enriched for glial cell type markers, oligodendrocytes (BW-M7), shows an enrichment for LoF-intolerant genes, but its degree of enrichment is far lower than that of neurons (**table S2**).

Consistent with prior work linking LoF intolerant genes with neurodevelopmental pathways, the most strongly-enriched module, BW-M1, enriches for markers of neural progenitors,^50^ neuronal migration,^51^ neuronal differentiation, and neurogenesis.^52^ This module is most strongly preserved in known neurogenic regions (such as the hippocampus **figure 1d**), suggesting that it corresponds to neural progenitor cells (NPCs) and adult neurogenesis. BW-M5 (MAPK cascade, ATP processing) enriches for the KEGG pathways of neurodegenerative disease: Huntington’s, Alzheimer’s and Parkinson’s diseases (**figure S2c**). The concentration of LoF-intolerance within neuron and neuron-related modules suggests either a cellular or a tissue-level buffering for genetic disruption to microglia and astrocyte specifically-expressed genes. In contrast, given that the function of oligodendrocytes is directly related the speed of neural transmission via ensheathment of axons, the enrichment in neurons and to a lesser extent oligodendrocytes suggests that efficient and rapid neural transmission could be genetically constrained.

To examine neuron-linked modules across regional hierarchies in greater detail, we use single-cell data^53,54,55^ aggregated by PsychENCODE to estimate cell type proportions among telencephalon samples via non-negative matrix factorization.^56^ By correlating sample cell type proportions with module loadings (eigengenes), we identify region-specific excitatory and inhibitory neuronal modules (**figure 2e**) including a Brodmann-specific (BA9+BA24) interneuron module M8, and an excitatory cell type Ex7 which loads only onto PFC-specific modules M1 and M3. Interestingly, no module corresponds identically to a single neuronal cluster, suggesting that either larger sample sizes or a more sophisticated gene-subclustering approach are required to achieve single-cell comparable resolution.

We observe that neither the striatal, nor cerebellar co-expression networks contain module sets corresponding to BW-M4 (neuron), suggesting that the neuronal subtypes in striatum and cerebellum give rise to sufficiently distinct co-expression patterns that form their own region-specific modules: CEREB-M2 represents cerebellar basket cells, CEREB-M3 represents Purkinje cells, and STR-M1 enriches for markers of medium spiny neurons MSNs, all of which are region specific cell types (**figure 2d**). To investigate this possibility, we expanded our previous approach to include single-cell sequencing from human cerebellum^57^ and mouse striatum,^58^ and identify three regional modules – BROD-M8, CEREB-M2, STR-M1 – with a strong relationship between module kME and relative gene expression in interneurons, Purkinje cells, and medium spiny neurons respectively (**figure 2c**). Consistent with the prior observation that constrained genes broadly enrich within neuronal genes, hub genes in these neuronal subtype modules are significantly enriched in those that are LoF-intolerant (**figure S2d**).

### Human-specific expression reflects differences in regional patterning

Recent comparative expression studies have identified thousands of genes up-regulated in humans compared to non-human primates, and have implicated spatial differences in neuronal subtypes and neurotransmitter receptors in driving this divergence.^59,60^ We sought to investigate whether patterns of human-specific regulation are reflected in regional biology, or whether the identified differences instead reflect inter-regional differences and brain patterning more broadly. To do this, we subset the co-expression modules from human and non-human primate brains identified by Sousa et al.^61^, to the 25 that show evidence of human-specific expression (**Methods**), 10 of which show significant overlap with our co-expression modules. Seven of these modules of human-specific regulation overlap whole-brain modules, and only two modules (M160, M220) solely overlap a regional module (**figure 2g**). These findings highlight that differential expression across regions and species generate different relationships from co-expression within regions. As such, regional differences in co-regulation between species remain largely unexplored.

### Identifying cell-type-specific lncRNA

Long non-coding RNA (lncRNA) are a diverse set of RNA species that modulate gene expression or protein function^62^ across many CNS cell types,^63^ and several studies suggest that lncRNA dysregulation is a component of neuropsychiatric disease.^64,65,67^ Many brain-expressed lncRNA have roles in neurodevelopment,^68^ and the enhancer with the most accelerated substitution rate in the human genome, HAR1, corresponds to the neuronally-expressed lncRNAs now termed HAR1A and HAR1B.^69^ Since lncRNA as a class tend to be expressed at a lower level than protein-coding RNA,^70^ they are difficult to profile and annotate through single-cell sequencing. We leveraged our co-expression networks corresponding to cell types to annotate human brain-expressed lncRNA in an unbiased, transcriptome-wide manner to associate lncRNA with neurological cell types and processes.

Only 52 known lncRNA species were profiled in the initial GTEx data set, likely because GTEx utilized library preparation using poly-A selection which would only profile the small subset of lncRNAs that are polyadenylated. Therefore, we expand the set of profiled lncRNA by projecting our whole-brain modules into ribosomal RNA depleted gene expression data from 44 neuro-typical post-mortem brains^71^ in which our whole-brain and cortical modules were well-preserved (ZSummary from 3 to 30; **methods**). We use gradient boosted trees^72^ to learn expression signatures of our module assignments in the new dataset and then classify lncRNA into their appropriate modules (**methods**). We identify 286 lncRNA belonging to major cell types and processes, the majority of which associate with neuronal module BW-M4 (66) or NPC module BW-M1 (109). Remarkably, slightly more than 20% (61/286) of these cell-type specific lncRNAs were previously shown to be dysregulated in neuropsychiatric disease (**table S4**). We cross-reference the inferred modules with published hippocampal and cortical RNA-seq,^73^ observing that lncRNA in our cell-type modules are indeed up-regulated within those cell types (**figure 2f**). Imputing Brodmann-area modules into the cortical RNAseq data identifies two predicted interneuron BROD-M8 lncRNA, *LINC000507* and *LINC00936* (RP11-591N1.1).

A previous study of ASD differential expression highlighted the differential expression of lncRNA as an integral component of the ASD transcriptomic signature.^74^ Because our module assignments for lncRNA are built from external data, we reasoned that we could use them to investigate differences in lncRNA co-regulation between ASD and control brain, and investigate whether lncRNA exhibit a different pattern of disruption in ASD from protein coding genes. We remove a set of matched protein coding genes (**methods**) from GTEx data, and performed module imputation for both lncRNA and matched protein-coding genes. After imputing gene modules using the neurotypical control samples, we contrasted both the expression and co-expression of module genes between ASD cases and matched controls. We find significant case-control differences in both expression and connectivity for modules BW-M1, BW-M6, and BW-M8 (p < 10^-15^, two-sample nonparametric Kolmogorov-Smirnov test), implying dysregulation in neurogenesis, astrocytes, and reactive glia. This observation is mirrored in the protein-coding genes, implying that lncRNA are disrupted in ASD to the same extent as other RNA species (**figure S2e,f**), as suggested previously.^75^

### Identifying cell-type-specific gene isoforms

Given the successful annotation of lncRNAs using co-expression, we apply a similar strategy to integrate isoform-level expression within cell type modules, both to understand cell-type specific splicing and to identify those isoforms likely involved in specific pathways (**figure 3a**). We identify 1,987 isoforms showing specificity to major cell types (**methods**) as measured by isoform expression (transcripts per million) kME to cell type modules, of which 549 are neuron, 543 astrocyte, and 696 oligodendrocyte. To validate these findings, we obtain RNA-sequencing data from sorted cells,^76^ quantify expression at the isoform-level, and rank isoforms and isoform ratios within cell type (**methods**). As expected, we find a significant association between isoform kME (to a cell-type module) and the rank of that gene’s expression in the sorted cell data (Spearman’s rho=0.286 oligo 0.258 astro, p<10^-15^ for both, **figure 3b**).

**Figure 3:**
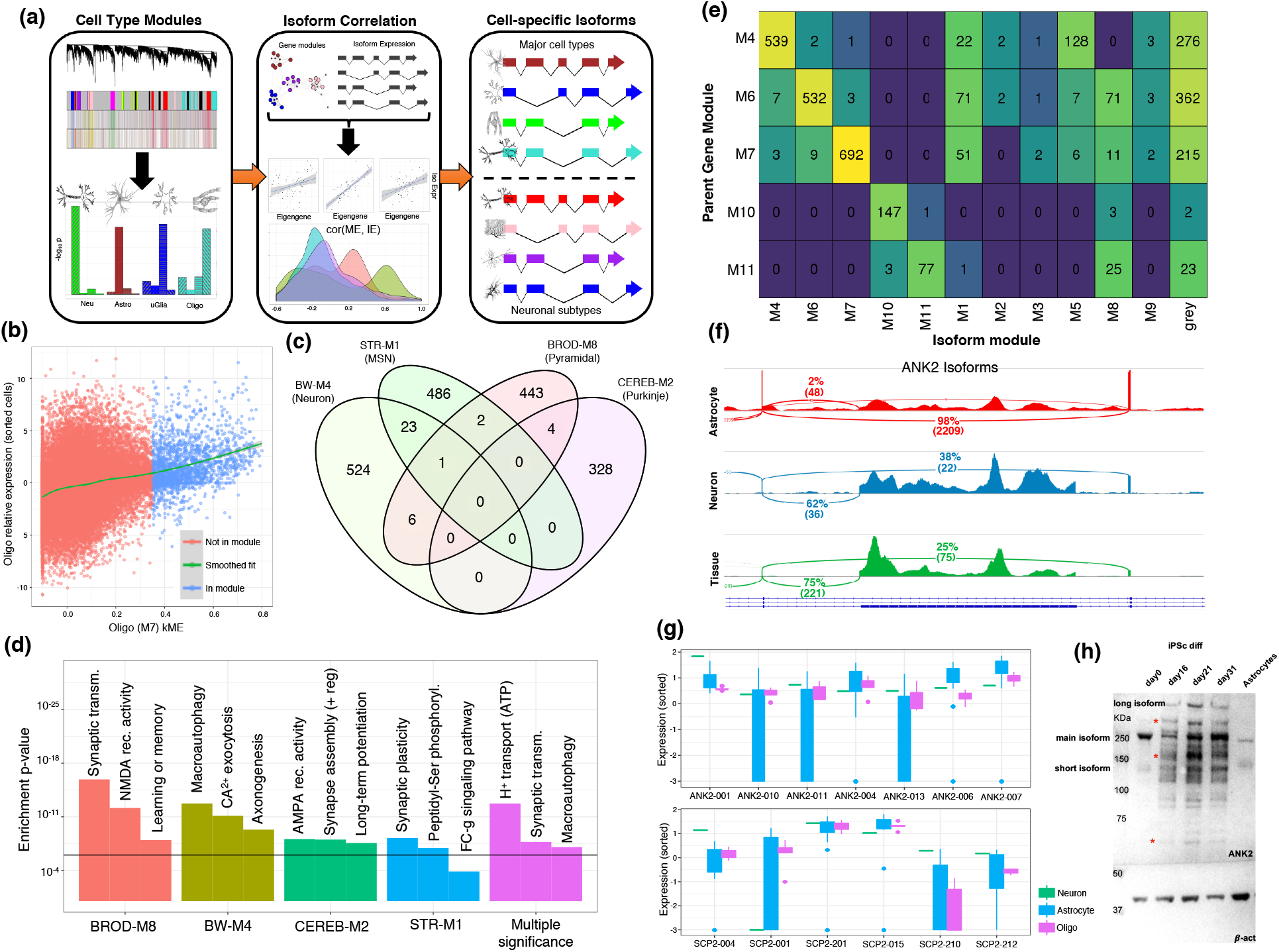
Creating a catalogue of cell-specific isoforms. (a) Overview of isoform assignment on the basis of kME to cell-type modules. (b) Isoform relative expression (log-FC of TPM) in oligodendrocytes plotted against isoform kME to BW.M7 showing significant positive relationship (p<10-6, linear regression). (c) Venn diagram of isoforms assigned to neuronal subtypes (d) GO enrichment of parent genes of subtype-specific isoforms. Top module-specific terms are shown, followed by terms which are significant across multiple subtypes (min p-value shown). (e) Assignment of daughter isoforms of genes with membership to a whole-brain cell type module, showing that most daughter isoforms are either assigned to the parent gene module, or to the grey (unclustered) module. (f) IGV visualization of the event differentiating the astrocyte and neuron isoforms of ANK2, the inclusion of the giant exon, in sorted cell data. (g) Expression of ANK2 and SCP2 transcripts in sorted-cell data, showing isoform switching between neuron and astrocyte. (h) Western blot of ANK2 across iPSC differentiation into neurons, and within astrocytes, demonstrating the presence of a long isoform specific to neurons.

Since single cell data does not yet provide similar isoform level coverage to bulk data, this validation of our approach motivates us to build putative cell-specific isoform maps for the region-specific cell types for which we had an enriched module: D1/D2 medium spiny neurons, Purkinje cells, basket cells, and inhibitory neurons. We identify between 300 and 500 putative cell-type-specific isoforms, finding that very few (<5%) of the resulting isoforms show high kME to the broad neuronal module BW-M4, consistent with the interpretation that these marker isoforms are cell type specific (**figure 3c**). All isoforms enrich for synapse-related functions, with additional regional variability: MSN isoforms enrich for the oxytocin signaling pathway, consistent with the observation that they are downstream targets of oxytocin in the nucleus accumbens,^77^ whereas Purkinje isoforms enrich for AMPA receptor regulators, and inhibitory neuron isoforms enrich for the NMDA receptor activity (**figure 3d**). These results demonstrate how our networks can be used to identify cell-type-specific splicing differences from bulk expression data, and highlight the synapse as a nexus of gene regulatory complexity and heterogeneity. Furthermore, they provide further evidence for the importance of isoform level analysis compared with gene expression alone in defining cell-type specific transcriptomes.^78^

### A subset of ASD risk genes switch isoforms across cell types

We next leverage this rich cell type-specific isoform data to identify examples of single genes with different isoform profiles across distinct cell types (“cell switching”). In the vast majority of cases the parent gene and cell-specific isoforms are assigned (by virtue of high kME) to the same co-expression module. However, in 7% of cases, the parent gene of an isoform will confidently differ in co-expression relationships from at least one of its alternatively spliced derivatives (**figure 3e**). We identify 52 genes exhibiting isoform switching between cell type modules, 11 of which show neuron/astrocyte switching (BW-M4/BW-M6), and 8 or which show neuron/oligodendrocyte switching (BW-M4/BW-M7). In all cases, the expression trend from sorted-cell RNA-seq matches the kME trend observed between isoforms and cell-type specific modules (**figure S3a**). Notably, of the 11 neuron/astrocyte switching genes, two, *ANK2* and *SCP2*, are well known autism spectrum disorder (ASD) susceptibility genes (**figure 3f**), while two others, *ERGIC3* and *PDE4DIP* are strong candidates (AutDB^79^ score 4; **figure S3b-e**; p < 0.01, Fisher’s exact test for enrichment of AutDB>4 genes among neuron/astrocyte switching genes). *ANK2* and *SCP2* show differential splicing of at least one event in ASD vs CTL brains (Parikshak2016 data, FDR < 0.05, linear mixed-effects model), but *ERGIC3* and *PDE4DIP* do not, and *ANK2* isoform switching is exhibited between ASD and SCZ cases.^80^ Analysis of single-cell data indicates that the primary difference between *ANK2* neuron- and astrocyte-specific transcripts is the inclusion of the 2,085 amino acid “giant exon.” The giant exon of ankyrins are known to function as an organizer of initial axon segments, and as a stabilizer of GABA-A synapses,^81^ confirming a neuron-specific role for this splice variant of *ANK2*. We further validate this finding at the protein level by western blot at several maturation points (day 0, day 16, day 21 and day 31) of iPSC differentiation into neurons and astrocytes, establishing that the long isoform is absent in progenitors and astrocytes, but present and persistent in neurons (**figure 3h**), suggesting that the large exon of *ANK2* is of particular relevance to some of its neuron-specific functions. However, the differentially spliced events for *ANK2* in ASD and SCZ do not involve the giant exon, indicating that while there is a role for the disruption cell-type specific alternative splicing in ASD, it is not related to up-regulation of this glial isoform within neurons.

### Ribosomal genes are down-regulated across the cortex

We next sought to incorporate regional differential expression with co-expression to provide insight into module-level regulation across brain regions. While differential expression has been used to identify differences between brain regions^82^, we find that this approach is too broad: nearly every gene expressed in brain shows differences in expression across brain regions (n=15616/15894, FDR<10^-3^, likelihood ratio test). We reasoned that genes with “extreme” expression profiles (i.e., significantly up-regulated or down-regulated in a region-specific or multi-region-specific manner) are more likely to have a specific role. We developed a novel Regional Contrast Test (RCT, **figure 4a**), which assigns z-score per group of regions per gene, whereby z-scores > 4 represent over-expression within that group, and those with z-scores <-4 represent under-expressed (**methods; table S5**). Because of the presence of several similar regions, we examined contrasts at varying degrees of granularity (**table S2**).

**Figure 4:**
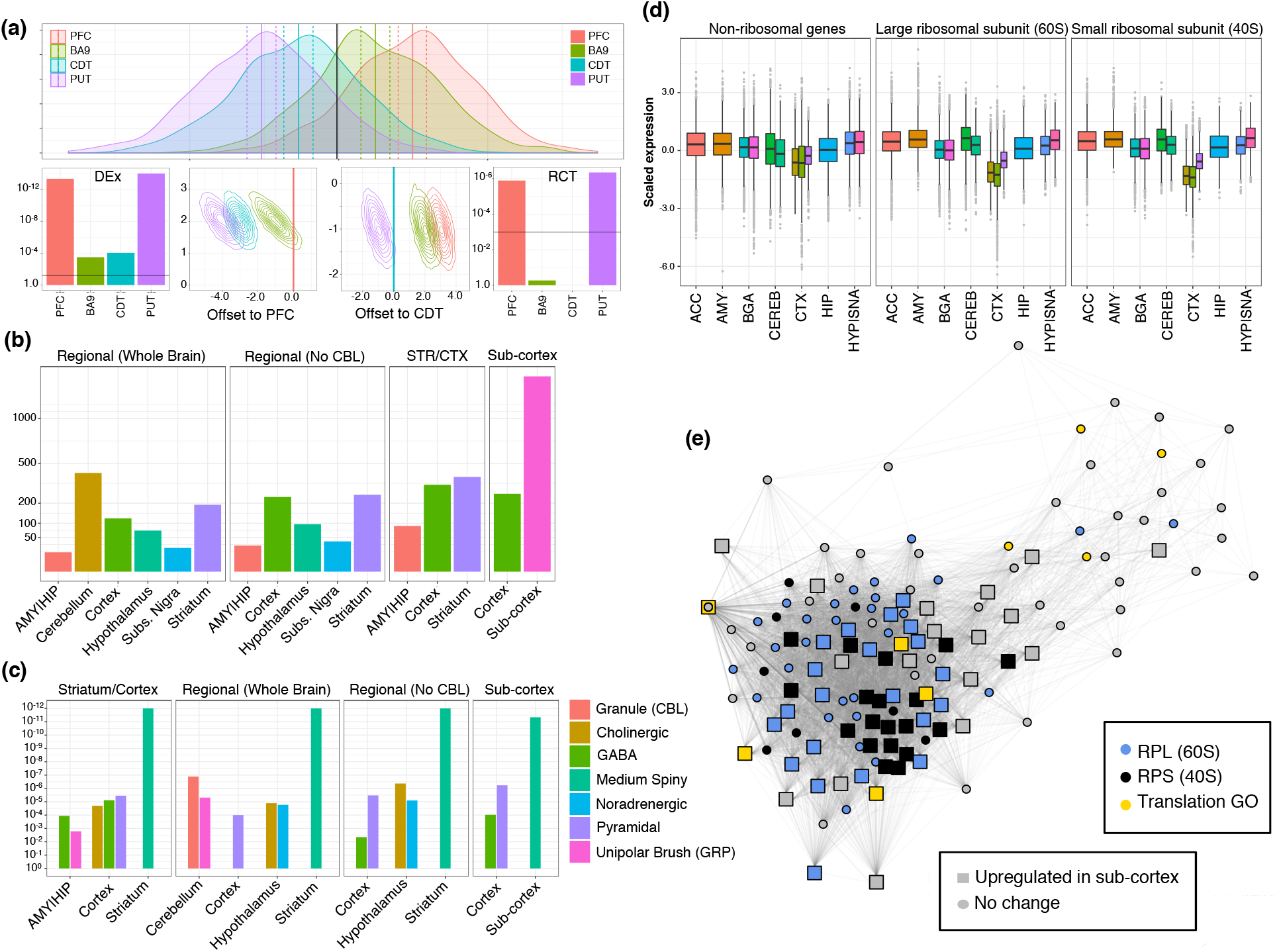
Region-specific gene up-regulation reflects region-specific cell types and ribosomal turnover. (a) Overview of the regional contrast test: top Example of heterogeneous data where mean expression within each region differs from the global mean. left With only 50 samples, all regions are significantly differentially expressed in a global manner. middle Visualization of the PFC statistic for PFC and CDT: The PFC mean (set to 0) overlaps only a small amount of the confidence region for another region, while confidence regions straddle the CDT mean. right The RCT statistic identifies the two most extreme tissues as differentially up- and down-expressed compared to all other regions. (b) Count of genes, which are significantly up-regulated within brain regions, across four backgrounds. (c) Cell-type enrichments for the up-regulated genes for the corresponding comparisons in (b). (d) Plot of scaled expression (per gene across tissues) for all genes in BW-M2, showing CTX-specific down-regulation of ribosomal subunits. (e) PPI co-expression network for genes in WB-M2, showing a sizeable fraction of the module core, a substantial fraction RPL and nearly all RPS mRNA are up-regulated in sub-cortical regions.

We first examine the set of genes up-regulated in subcortical regions (striatum, hippocampus, and amygdala) versus cortex, and observe that these differences enriched for non-neuronal cell type modules (p < 1e-10 for BW-M11, BW-M6, BW-M8, BW-M10, and BW-M7), consistent with a higher glia/neuron ratio in the striatum (**figure 4c**). Conversely, we find BW-M4 (neuronal) to enrich for the genes up-regulated in cortex, as compared with sub-cortical regions. Interestingly, we also observe a significant (p = 4.89e-3) enrichment in BW-M2, a module dominated by small- and large-ribosomal subunit RNA (**figure 4d, 4e**), for sub-cortical upregulated genes. This suggests that subcortical regions may show higher translational demand, more ribosomes, or faster ribosome turnover than cortical regions. Recent work has demonstrated that protein turnover – particularly the large and small ribosomal subunits – drastically increases in cultures with high glial proportion,^83^ providing an explanation for increased abundances of these ribosomal mRNA in regions of high-glial-proportion in the brain.

### Qualifying regional specificity of previously-identified neuropsychiatric disorder co-expression networks

We next use this multi-region data set to address regional specificity of disease associated modules by re-evaluating gene modules identified in post mortem tissue from 11 publications (normal brain^84,85^, autism^86^, schizophrenia^87,88^, cross-psychiatric^89^, Alzheimer’s disease^90^, epilepsy^91^, and developing brain^92,93,94^) with the objective of identifying: i) whether or not those previously-discovered modules, typically generated using only one or two brain regions were related to any the modules we identified from the normal individuals in GTEx, and ii) whether those previously-discovered modules are indeed region-specific. Remarkably, we observe a common set of modules involved in overlaps across every study: BW-M1, BW-M4, BW-M6, and CTX-M1 (**figure S4a-d**). In fact, there are no studies for which a disease-implicated module does not show significant overlap with one of the GTEx whole-brain or multi-regional modules, suggesting that these many studies likely reflect overlapping co-expression patterns corresponding to core cell types disrupted by neuropsychiatric disease (**table S6**).

### Convergence of molecular signatures into brain-wide and regional pathways in neuropsychiatric disease

We next investigate whether genetic perturbations in neuropsychiatric disease converge onto region-specific or cross-regional modules. Utilizing databases of *de-novo* variants implicated in ASD and SCZ,^95^ GWAS summary statistics,^96,97,98,99^ and our RNA-sequencing in post-mortem ASD and normal brains,^100^ we identify two whole-brain modules, BW-M4 (neuron) and BW-M1 (neural progenitor) that simultaneously enrich for ASD-linked rare variants (**figure 5a**), SCZ GWAS signal (**figure 5b**), and that manifest disrupted expression in ASD post mortem brain relative to controls (**figure 5c-g**). We also identify two regional modules, CTX-M3 (activity-dependent regulation and endocytosis) and CEREB-M1 (mRNA binding) that show ASD rare-variant and SCZ GWAS enrichment. While the co-expression relationships for CTX-M3 and CEREB-M1 are distinct, the genes do overlap more often than expected by chance (Jaccard=383/1938, OR=9.5, p<10^-20^ Fisher’s exact test), and both show significant preservation in control brain, but not in ASD post mortem brain (**figure 5g**), consistent with the disruption of these modules in ASD.

**Figure 5:**
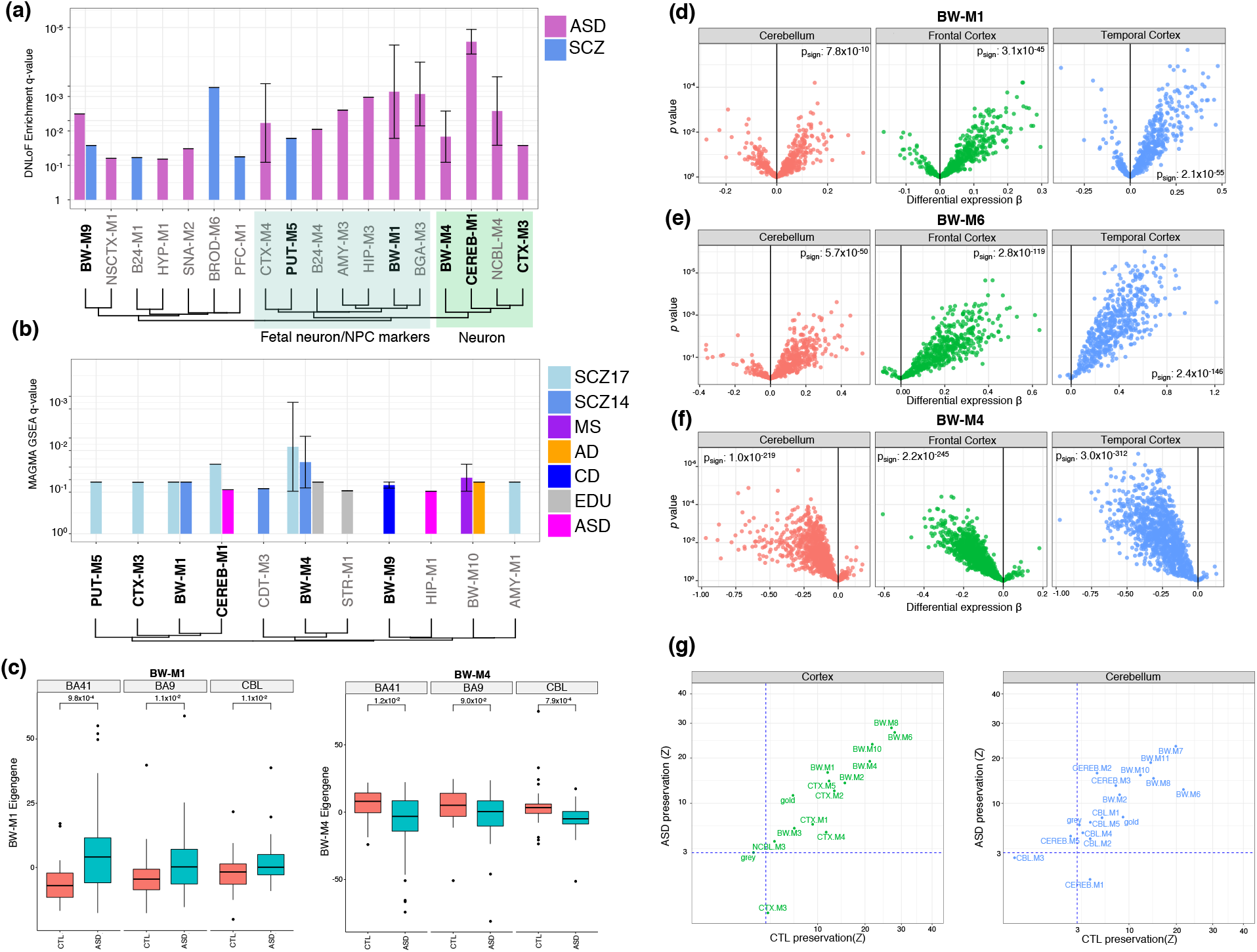
Gene-level module enrichments for de novo PTVs, GWAS summary statistics, and differential expression. (a) FDR values (Fisher’s exact test) for enrichment of de novo loss-of-function variants within modules, summarized to module sets. Bar height gives geometric mean of FDR, and whiskers the range of (significant) FDR values for modules within the module set. Bold modules also show enrichment from GWAS summary statistics; see (b). Module sets are ordered by Jaccard similarity between their index modules. Green region: These modules enrich for neuronal markers. Blue region: These modules enrich for fetal neuron, mitotic progenitor, or outer radial glia markers. (b) FDR values (MAGMA) for GWAS summary statistics within modules. Method of ordering identical to (a). (c) Module eigengene expression for BW-M1 and BW-M4 in ASD cases and control brains across three regions and associated p-values from a T-statistic (linear model including covariates as in Parikshak2016). (d-f) Volcano plots and sign-test P-values for genes in NPC, astrocyte, and neuronal modules. (g) Module preservation statistics separately in ASD and control brains, suggesting differential preservation for modules CTX-M3 and CEREB-M1.

BW-M4 represents a neuronal module set, identified independently throughout the telencephalon and sub cortical regions, all sharing GO terms related to membrane organization or ion transport (**figure S5a**). Examination of significant genes within BW-M4 (defined as MAGMA Z>3.0, **Methods**), we find that the enriched terms for both the Psychiatric Genomics Consortium^101^ and ClozUK^102^ SCZ GWAS studies are related to the synapse and synaptic transmission (**figure S5b**), suggesting a convergence of risk genes onto synaptic signaling pathways, consistent with a recent comprehensive pathway analysis.^103^ Using meta-GSEA (**methods**) to rank ontologies across GWAS studies and brain regions, we find that both ASD and SCZ appear to share highly-ranked terms related to synapse assembly and plasticity. The term synaptic transmission shows very strong evidence only from SCZ association statistics, while the strongest terms with evidence in ASD alone are learning and social behavior (**figure S5c**). Because of the difference in GWAS power, it is likely that ASD-specific terms reflect ASD-specific biology: ASD is a social disorder that can present with learning disability. On the other hand, the implication of synaptic transmission in SCZ alone likely reflects that sample sizes are too small, and thus power too low, to draw mechanistic conclusions about ASD genetic risk.

BW-M1 contains genes and pathways corresponding to neurogenesis, differentiation, and migration (**figure 6a**), as well as components for RNA splicing, structural components of cell division, and stem cell population maintenance (**figure 6b**). Genes within BW-M1 are strongly loss-of-function intolerant, and the module enriches significantly for PPI interactions. The genes in this module, which are up-regulated in ASD cortex, include the TGF-beta signaling pathway (FDR=0.0047, STRING^104^). This pathway is known to regulate neurogenesis,^105^ and consist mainly of the BMP/SMAD pathway (*BMPR1A*, *BMP2K*, *SMAD4*, *SMAD5*, *SMAD9*) which is critical for orchestrating proliferation/differentiation balance.^106^ NPC proliferation/differentiation balance is another major theme of the module, as it contains key *REST* co-repressors *CTDSPL* and *RCOR1*, the down-regulation of which promote proliferation over differentiation;^107^ as well as differentiation repressors *ADH5*, *TLR3*, *SOX5*, *SOX6*, and *PROS1*^108,109,110,111,112,113^ and the differentiation/proliferation regulator *SPRED1*.^114^ The *de novo* LoF and GWAS enrichments suggest that NPC proliferation and differentiation – both in the prenatal and adult brain – are disrupted in ASD and SCZ. Analysis of module trajectories in the developing brain^115^ shows very strong prenatal upregulation, with continuing postnatal expression into early adulthood (**figure 6d,e**). Disruption of this module may be related to the observed ASD expression signature – downregulation of neuronal modules and upregulation of astrocyte modules^116^ – suggesting that one component of neuropsychiatric disorders are brain-wide changes in neuronal proliferation/differentiation/maturation balance that begins in early development and persists throughout postnatal development and into adolescence.^117^

**Figure 6:**
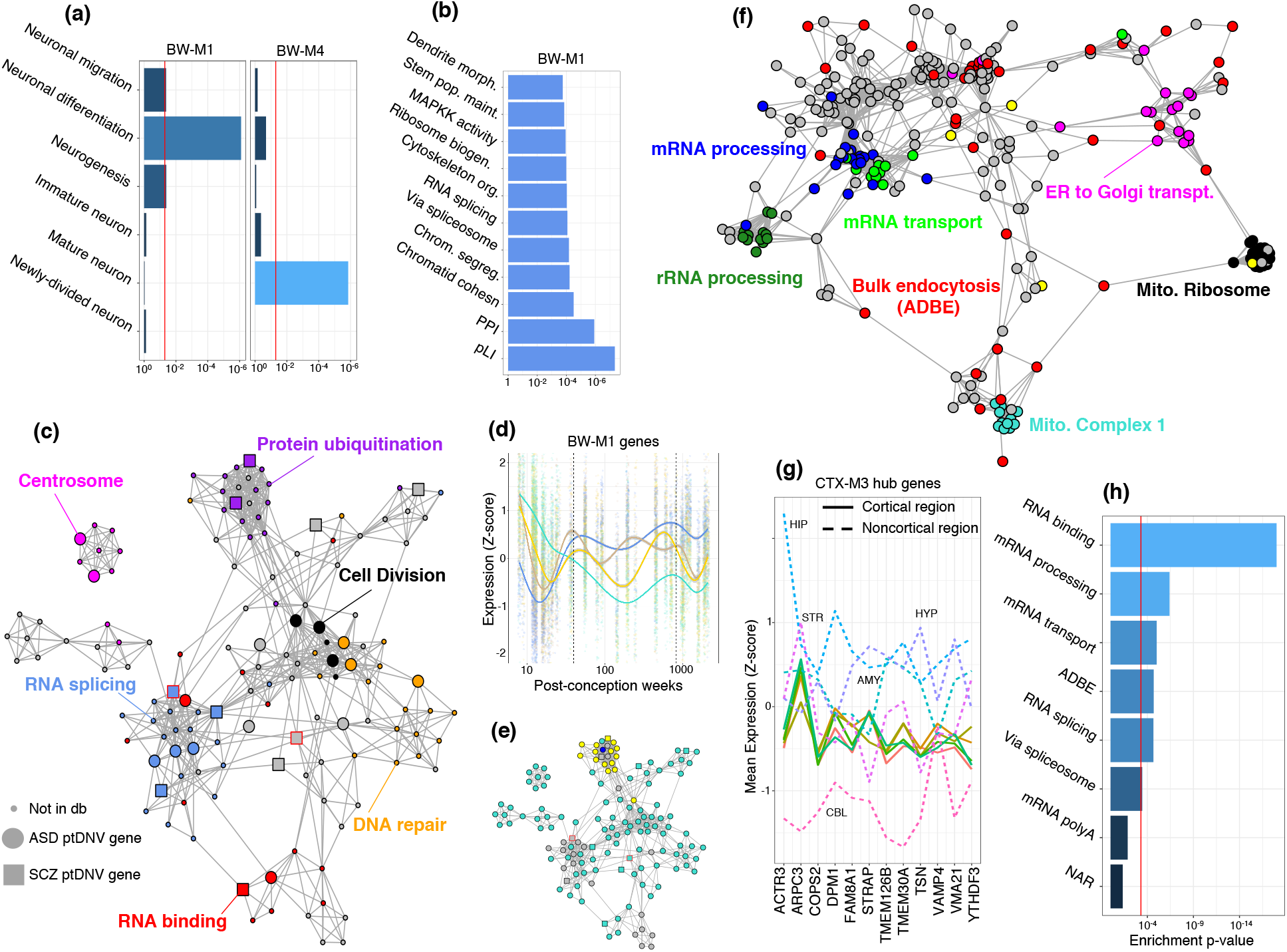
Ontologies, PPI networks, and expression profiles of ASD-associated modules. (a) Enrichment p-values (Fisher exact test) for neuron-related ontologies in whole-brain modules. (b) Combined (geometric mean) enrichment p-values of ontologies for all modules in module set BW-M1 that showed enrichment for ASD-implicated de novo loss of function mutations. (c) Co-expression-PPI network of BW-M1, highlighting de novo loss of function mutations (large nodes) and ontologies (colors). (d) Expression of BW-M1 across developmental time-points, subclustered into four modules using WGCNA. (e) Assignment of network nodes in (c) to the subclusters in (d) via label propagation. (f) Coexpression-PPI network for CTX-M3, colored by enriched gene ontology sets. (g) Expression profile of CTX-M3 hub genes across brain regions, demonstrating tight co-regulation in cortical regions (solid lines) by virtue of small variance, and variable expression across non-cortical regions (dashed lines). (h) Enrichment p-values (Fisher exact test) of the CTX-M3 module for gene ontologies, including bulk endocytosis genes.

The two regional modules that show convergent evidence of disruption in neuropsychiatric disorders, CTX-M3 and CEREB-M1, show an enrichment for *de novo* LoF variants linked to ASD, gene set enrichment for SCZ GWAS risk variants, and are disrupted in post mortem brain from ASD subjects (**figure 5**). Although, they have distinct components, both CTX-M3 and CEREB-M1 show significant overlap in their genes, and modest evidence of preservation outside of their respective regions, with preservation AUPR scores < 0.5 (**figure S6a,b**). Both modules enrich for PPI (CEREB-M1 p<7e-15, CTX-M3 p<0.0023), as well as LoF-intolerant genes, indicating that they contain essential biological pathways.

To validate the cortical-specificity of CTX-M3, we used normalized RNA expression values from the Allen Human Brain Atlas^118^ to contrast the expression trajectories of hub genes in cortical versus noncortical regions. We find that the relative levels of these genes across all cortical regions (frontal, parietal, occipital, and temporal lobes plus cingulate gyrus) are tightly coupled, in contrast to highly variable expression in non-cortical regions (hippocampus, hypothalamus, striatum, cerebellum, **figure 6g**), confirming cortex-specific co-expression.

Notably, CTX-M3 contains both the syndromic ASD risk gene, *FMR1*, as well as its direct interactor, the protein *NUFIP1*, both of which have been implicated in the regulation of activity-dependent translation and local synaptic translation^119^ and additionally in ribophagy.^120^ It also contains the ID gene *ATRX*, which forms a complex with the protein product of *DAXX* to regulate H3.3 loading onto and maintenance within heterochromatin. H3.3 is itself associated with activity-dependent transcription in neurons,^121^ suggesting that dysfunction or dysregulation of *ATRX* could alter the availability of this activity-related histone.

To further examine the role of activity-dependence within this module, we examine a set of genes identified as up-regulated following neuronal activity,^122^ finding that 10% of these genes fall into CTX-M3 (p = 0.0472, Fisher’s exact test, **figure 6h**). The observed enrichment is driven largely by components of protein phosphatase 1, which has both nuclear and synaptic roles in synaptic plasticity and long-term memory.^123,124^ However, several components of the mitochondrial ribosome (*MRPL27*, *MRPL45*, *MRPS26*) are observed concomitant with activity-dependent upregulation, and CTX-M3 is highly enriched for the mitochondrial ribosome, containing 21 genes within this functional pathway (p < 1.7e-10, Fisher’s exact test). These observations indicate that genes normally up-regulated by neuronal activity form one component of this down-regulated ASD and SCZ associated module, CTX-M3. Other components of this module include poly-A binding, alternative polyadenylation and alternative splicing (*NGDN, MBNL1, MBNL2, CSTF3, SPSF3, CPSF6*), multiple endocytosis regulating genes (*RALA, VAMP4, VAMP7, TSG101, VPS25, RAB18, RAB3GAP2, CHMP2B*, and sorting nexins *SNX2, SNX3, SNX13, SNX14*), consistent with their potential role in supporting neuronal activity dependent processes that are disrupted in ASD (**figure 6f**).

### Network enrichment and genetic architecture: quantifying omnigenics in neuropsychatric disease

Complex diseases including neuropsychiatric disorders are influenced by large numbers of genetic variants and genes.^125,126,127^ Gene networks from disease-relevant tissues can capture interactions between these genes and have therefore been hypothesized to inform disease heritability.^128,129,130^ Gene-set enrichment analysis of network modules is a simple approach for investigating how network structure relates to disease-associated gene. However, this approach ignores network distance and higher-order structures and does not directly inform models of heritability. We therefore sought to evaluate the use of network distance, as defined by brain-wide and regional co-expression networks, into a model of genetic architecture, and examine the role that co-expression networks play in the genetic architecture of neuropsychiatric disease.

Our approach is motivated by the recently proposed omnigenic model of disease.^131^ The omnigenic model posits the existence of core and peripheral genes: the core genes are defined as those with direct effects on disease, with the peripheral genes manifesting effects that are mediated through the core genes. That is, within a cell type or set of cell types relevant to disease, mutations in peripheral genes are proposed to lead to increased susceptibility due to indirect perturbations of core gene activity. This model suggests that larger disease effect sizes will be observed for loss-of-function variants affecting core genes, common cis-eQTLs of core genes, and variants affecting peripheral master regulators that converge on numerous core genes.^132^ Embedded within this formulation is the expectation of two broad classes of genes: one class of genes with high effect size mutations (core genes and peripheral master regulators), and another class with small indirect effects that act in *trans* (peripheral genes^133^). Although the omnigenic model does not explicitly define how core and peripheral disease genes relate to gene networks, co-expression networks naturally exhibit a hub and periphery structure, and it has been hypothesized that module hub genes are more likely to harbor large effect size mutations,^134^ and thus may be potential candidates for core disease genes or their master regulators. We therefore sought to test whether network-central genes in brain co-expression networks might reflect an omnigenic disease architecture.

With this motivation, we first construct a simple model whereby allelic effect size is a function of network distance to simulated core genes (network genetic architecture, **methods**). We simulated variants to generate a frequency-effect-distance distribution (**methods**), and observed that the resulting effect size and heritability distributions bore a striking resemblance to the theoretical distributions posed in the original omnigenic publication (**figure 7a**).^135^ We observe that within this model, high-effect variants fall within or very near to core genes, and nearly all of (98-100%) the top 1% of variants by effect magnitude fall into the first decile of distance to the simulated core genes (**figure 7b**). When the distance is corrupted by 30% random measurement error (**methods**) this proportion remains robustly above 60% (**figure 7b**).

**Figure 7:**
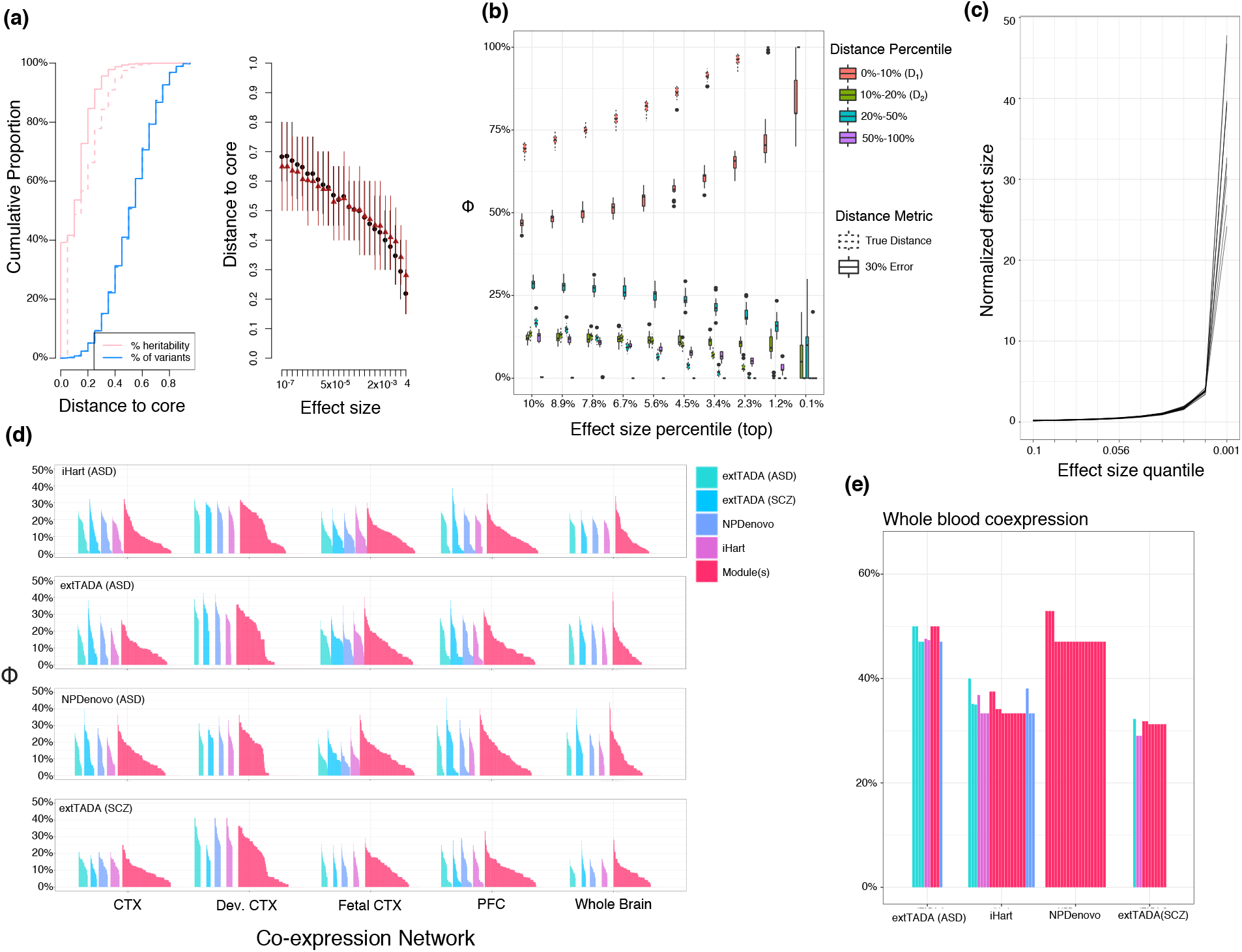
Characterizing core-periphery structure of high-impact neuropsychiatric disease genes across multiple networks. (a) Example simulation of network genetic architecture, where the variant effect size decays rapidly with distance to core. Left: Cumulative proportion of genes (blue) and heritability (pink) along the distance distribution. Dotted line shows the cumulative heritability when true distance is replaced by a corrupted (30% error) distance. Right: The relationship between core distance and effect size results in high-effect variants only appearing very close to core genes. (b) High-impact genes are defined by the effect-size percentile on the x-axis, and the % of genes falling into the core-distance decile is plotted on the y-axis. This plot encompasses 20 simulations. Dotted boxes represent the expected values for Φ when the distance is error-free, while solid boxes represents the case where distance is 30% corrupted by error. (c) Validation of the effect size distribution: the effect size of each quantile is normalized to the effect size for which a balanced GWAS of 10,000 samples has 80% power; the highest-impact variants are only 20-50x stronger than empowered variants. (d) All values of Φ across distance metrics, core set size, module definitions, and brain co-expression networks, demonstrating that no value of Φ exceeds 50%. (e) top 10 Φ values (per core set) for the GTEx whole-blood co-expression network.

We leveraged our robust, comprehensive brain networks to evaluate the extent to which various gene networks’ central genes capture an omnigenic “core” structure for two common neuropsychiatric disorders: ASD and SCZ.^136^ We reasoned that genes associated with these disorders by virtue of *de novo* loss-of-function mutations are likely to be both: (i) core genes under an omnigenic architecture, and (ii) among the highest-effect mutations for these diseases (core genes under our model). We identified sets of genes likely to harbor high-impact rare variation for both ASD and SCZ by using the top implicated genes from each of three previous rare and *de novo* studies of neuropsychiatric disease: extTADA,^137^ iHART,^138^ and NPDenovo^139^ (**methods**). We leveraged these data to develop a statistic, *ϕ̂*_core_, that measures the “core” nature of a hypothesized gene set on an arbitrary network. We then use this approach along with brain co-expression network structure to evaluate whether either 1) network-central genes or 2) rare-variant implicated genes appear to behave as “core” genes (**methods**). Specifically, we define the core proportion of a network, *ϕ̂*_core_, as the proportion of likely high-impact genes whose distance to proposed core genes (e.g., network central genes) is in the lowest decile. For instance, using 10 genes as a proposed core and a 10,000-gene network results in 10,000 distance values (minimum path distance to any of the 10 genes), *ϕ̂*_core_ is then the proportion of rare-variant implicated genes whose rank (by distance) is 1,000 or less. A value of *ϕ̂*_core_=1.0 would occur if all true disease core genes fall very close to the proposed core genes within a network (e.g., if disease core genes are network central genes) reflecting a well-separated core/periphery structure.

First, in an exploratory analysis, we evaluated network-central genes across whole brain, cortex, prefrontal cortex, developing cortex, and fetal cortex co-expression networks, using whole blood co-expression as a baseline. We used multiple approaches to selecting modules and hub genes, for defining co-expression distance, and for selecting the number of rare-variant implicated genes (**methods**), resulting in 6,732 statistics for brain tissue networks and 7,124 statistics for the whole blood network.

As expected, the largest observed value of *ϕ̂*_core_ from whole blood co-expression networks using network central genes was 0.52, below the simulated theoretical baseline (**figure 7**). Surprisingly, the largest observed value of *ϕ̂*_core_ across all cortical tissue co-expression networks and their central genes was even lower: 0.44, representing the distance of NPDenovo-implicated genes to module BW-M7 (**figure 7**), while other networks in our collection showed peak values ranging from 0.25 (whole cortex evaluated with extTADA-SCZ) to 0.43 (whole brain evaluated with extTADA-ASD). The fact that the statistic is higher for blood may result from the larger number of whole-blood modules, which increases the number of comparisons, or by the larger number of whole-blood samples which increases the power to separate and define modules. Importantly, *ϕ̂*_core_ does show a significant odds ratio for 416/6732 brain network distances (FDR < 0.05, Fisher’s exact test) and for 217/7125 blood network distances: values of *ϕ̂*_core_ as low as 0.207 are statistically significant, demonstrating that the genetic architecture of these diseases indeed reflect network distances, just not so strongly as to cleanly separate core and peripheral genes.

Because disease core genes need not be network-central genes, it could be the case that co-expression networks capture the correct notion of gene-gene distance, but network-central genes are not the correct core gene set. Therefore, we next evaluated use of rare-variant implicated genes as candidate core genes. We partitioned the rare-variant implicated genes from the source studies into proposal (10 or 20 genes) and evaluation sets (35 to 100 genes, **methods**), using the proposal gene set to compute network distances, and the evaluation gene set to calculate *ϕ̂*_core_. This resulted in 3,562 statistics for all brain co-expression networks, and 484 for the blood co-expression network. As in the network-central gene analysis, we find that the largest value of *ϕ̂*_core_ in brain networks (0.47) is lower than that of blood networks (0.50) which is in turn lower than the theoretical baseline of 0.60.

The inability of the co-expression networks to separate a core and periphery according to our test may reflect one of several explanations. First, it may be that the diseases assessed do not in fact have a core/peripheral gene structure, and the omnigenic model is insufficient to explain their architecture.^140^ Second, it could be that peripheral master regulators are somewhat common among the candidate core genes (e.g. *de novo* LoF genes) tested above, and thus not a “clean” core set. We have endeavored to reduce the impact of potential master regulators by excluding known transcription factors, DNA binding proteins, RNA binding proteins, and noncoding genes during our analysis (**methods**), so any master regulators would need to function through an alternative indirect mechanism. A third possible explanation is that the networks here are simply not the correct networks to describe the architecture of these diseases, whether due to power, cell type specificity, or other properties – we return to this possibility below. Finally, it could be that network centrality is not the correct property to consider in assessing omnigenic architecture, and that an alternative analysis using arbitrary connectivity without candidate network central genes would be required. While we cannot definitively assert which influences our results, it is clear from our analysis that the simple interpretation that co-expression network central genes will cleanly distinguish large effect “core” genes from a periphery is not supported.

To consider the impact of network choice on these results, we explored several possibilities. Although most modules identified in co-expression networks from bulk tissue do represent cell types assayed via single-cell expression,^141^ it is possible that either the bulk nature of tissue data, or the fact that these co-expression networks are based on static correlation, rather than physical or dynamical interactions, fail to capture the appropriate core-periphery relationships. To consider other types of networks, we utilized the InWeb PPI network^142^ from brain, and inferred empirical gene regulatory networks (eGRNs) from brain tissue and cell types (**methods**),^143^ and repeated the analyses above using high-connectivity genes as network central genes. We find that the core/periphery structure in these other networks also do not mirror the expectations of our omnigenic-like model (**supplemental figure 8**): across both PPI and eGRN networks, the largest observed value of *ϕ̂*_core_ was 0.38.

## Discussion

Gene co-expression networks have provided a powerful organizing framework for transcriptomic studies of the nervous system. Many gene co-expression network studies have focused on a single brain region, albeit relative to disease states, or over development.^144,145,146,147,148,149,150,151,152,153^ In many cases, this has made it unclear whether such networks were specific to the brain regions studied, or more generalizable across regions. Here, we have described the construction of a robust, hierarchical, resource aimed at establishing common and region-specific aspects of gene co-expression within the brain. We identified 11 whole-brain co-expression modules, corresponding to common cellular components such as major neuron and glial types. We also captured region-specific signatures dominated by regional cell subtypes. By using a consistent framework that allows us to understand the relationship of modules identified at different scales, we demonstrated that: i) the convergent pathways in neuropsychiatric diseases such as autism and schizophrenia are primarily reflected in neuronal and neurogenesis pathways that are common across brain regions, rather than specific to a single region; ii) Disease risk in ASD and SCZ is enriched in these down-regulated neuronal and neurogenesis modules; several of these modules reflect down-regulation of activity dependent transcriptional programs; iii) cell-type-specific lncRNA and isoform co-regulation can be included these networks, and that isoform-level analysis is likely essential to interpret disease associations; Iv) brain RNA co-expression, PPI, and co-regulatory networks do not cleanly capture the dichotomous core/periphery structure proposed by the omnigenic model. It will be important to further explore cellular-level or other types of gene networks.^154^ For example, within-cell co-expression networks – which are difficult to estimate from bulk tissue or even single-cell expression – may more directly relate to disease etiology and thereby show more omnigenic-like network architecture parameters than the networks investigated here.

The fact that gene expression markers for major cell classes can be identified from bulk tissue co-expression is now well-established.^155,156,157^ The relationship between module membership and cell-type relative expression demonstrates that co-expression analysis is a valid method of marker discovery for both cell subtypes and isoforms. Indeed, isoform kME values for striatal modules M1 and M2 may represent the most salient information regarding both isoforms with high relative expression human medium spiny neurons. As sample sizes grow larger, we expect to identify new subdivisions of co-expression modules, corresponding to ever-finer distinctions between underlying cell types.

We show, through analyzing lncRNA from a separate collection of brains – sequenced in a different location and with different technology – that the modules we have constructed are applicable to other, smaller, RNA-sequencing datasets, and retain the very same cell-type signals. This approach based on gradient boosted trees is potentially very powerful: it allows every gene in a new expression dataset to be identified as potentially region-specific or cell-type specific, without requiring large sample sizes for co-expression analysis or samples from multiple regions.

A similar approach was used to create isoform networks, using guilt-by-association via co-expression to assign isoforms to modules, which in many cases are cell-type specific. This is important, as single cell sequencing approaches that capture full-length transcripts are expensive; and those that do not only incompletely represent isoform expression profiles.^158^ Here, we provide a first generation set of 1,987 cell-type specific isoforms for major cell classes in the brain of which 549 are neuron, 543 astrocyte, and 696 oligodendrocyte. Remarkably, several of these isoforms, including 4 ASD risk genes, manifest isoform switching between neurons and glia. One of these *ANK2*, has been recently described and validated as having different isoforms in neurons and glia, which manifest distinct PPI.^159^ As long-read sequencing matures and is applied at larger scale, more complete cell-type specific isoform networks can be constructed using this approach. Our data indicate that this will be of substantial value in understanding disease-relevant variation.

Our findings that ASD-linked dnLoF mutations as well as SCZ GWAS signal enrich in brain-wide neuronal and neurogenesis modules underscore previous findings linking both common and *de novo* variation to synaptic genes,^160,161^ neuronal genes,^162^ developmentally-expressed genes,^163^ and neurogenesis pathways.^164^ Even though cortical regions contain a much higher proportion of neurons than other brain regions, the pattern of enrichment is not cortex-specific, and enrichments in the BW-M1 and BW-M4 module sets are significant across the brain, implying likely widespread effects of these genetic risk variants on brain function.

The only region-specific modules with convergent evidence across disease and modality were CTX-M3 and CEREB-M1, which appear to reflect activity-dependent transcriptional profiles identified in previous studies. Indeed, *VAMP4* – present in CTX-M3 – is an essential molecule for activity-dependent bulk endocytosis (ADBE),^165^ and several module proteins (including RAB GTPases *RAB7a* and *RAB18*) overlap with the ADBE proteome.^166^ This suggests a parsimonious explanation that this module concerns the maintenance of organelles and proteins required for long-term neuronal activity, (i.e., mitostasis and ADBE proteostasis), through activity-dependent mRNA transcription and neuropil targeting.^167^ CTX-M3 does show suggestive evidence of preservation outside of cortical regions (**figure S6**), and it may appear cortex-specific simply because it is easier to uncover neuronal biology in regions with the highest proportion of neurons, such as the cortex. At the same time, the hub genes of this module show tight regulation of mean expression across cortical regions, but not non-cortical regions in the Allen Human Brain Atlas,^168^ which supports weak, but not complete overlaps with other non-cortical neuronal modules. In addition, power to detect enrichment is a function of module size. Region-specific modules have a smaller average size than brain-wide modules, and it may be that additional samples will provide evidence for selective vulnerability as the number of samples increases.

Incorporating gene networks into models of genetic architecture remains a major challenge, with application both to predictive disease models and to translational genetics. Although the omnigenic hypothesis does not specify a concrete network model, in order to apply the model to etiology of specific diseases, we must connect it to quantifiable relationships between genes. Our approach to investigating the models comes from a unifying hypothesis: that there is a relationship between mutational effect size and network distance – with omnigenic and polygenic architectures representing the strong and weak extremes of that relationship. From this point of view, quantifying a network’s effect on trait architecture in terms of the decay parameter is of a higher concern than labeling it as either omnigenic or polygenic, as there likely are traits at both ends of this spectrum. For instance, one might expect secondary phenotypes of Mendelian disorders (such as age-at-onset or disease severity) to show a strong omnigenic-like relationship in the relevant network, with all distances measured to the disease gene. Our results show that organizing genes into bulk tissue co-expression networks does not explain high-impact mutations in terms of a clear core/periphery relationship – either from the perspective of modules, or empirically-defined clusters. For the three distinct network types tested – gene co-expression networks, PPI networks, and gene-regulatory networks derived from bulk tissue – we find that the network structures do not strongly distinguish peripheral genes from core genes as would be consistent with an omnigenic architecture. However, there are many other natural network topologies to test, including single-cell and within-cell networks, once sufficient data is available from a large number of subjects and cell types. Perhaps more importantly, the model underlying our analysis provides a means to relate total effect – direct and indirect – to a network structure. Future work in extending our model could provide a means of assessing the proportion of heritability explained by network interactions, and thereby probing the network architecture of disease.

## Expression quantification, QC, and covariate correction

Reads were aligned using STAR^clxix^ in standard two-pass fashion. Gencode v25 transcripts (hg19/b37) were used as the reference transcriptome and genome for alignment. Transcripts were quantified using RSEM to produce gene and isoform level TPMs. The analyzed TPMs are log-transformed log(0.005 + x) resulting in approximate normality.

Sample and individual-specific covariates were downloaded from the GTEx^clxx^ website, and supplemented with technical alignment information from the STAR alignment and PicardTools QC of the resulting .bams.

Individuals were excluded if they were positive for any of the following phenotypes: ‘MHALS’, ‘MHALZDMT’, ‘MHDMNTIA’, ‘MHENCEPHA’, ‘MHFLU’, ‘MHJAKOB’, ‘MHMS’, ‘MHPRKNSN’, ‘MHREYES’, ‘MHSCHZ’, ‘MHSEPSIS’, ‘MHDPRSSN’, ‘MHLUPUS’, ‘MHCVD’, ‘MHHIVCT’, ‘MHCANCERC’, ‘MHPNMIAB’, ‘MHPNMNIA’, ‘MHABNWBC’, ‘MHFVRU’, ‘MHPSBLDCLT’, ‘MHOPPINF’. The individual-specific covariates ‘GENDER’, ‘AGE’, ‘RACE’, ‘ETHNCTY’, ‘TRISCH’, ‘TRISCHD’, ‘DTHCODD’, ‘SMRIN’, ‘SMNABTCH’, ‘SMGEBTCH’, ‘SMTSISCH’, ‘SMTSPAX’ were extracted. The ‘DTHCODD’ variable was binned into the following categories: ‘UNKNOWN’, ‘0to2h’, ‘2hto10h’, ‘10hto3d’, ‘3dto3w’, ‘3wplus’.

STAR alignment metrics and PicardTools QC metrics were subset to non-excluded samples, outliers were flagged and removed via a chi-squared test (p < 10^-5^). The PicardTools metrics were log-scaled, and the top 5 principal components extracted using the PCA class from scikit-learn^clxxi^ (“seq-PC”). The STAR alignment covariates were subset to those with “splice” in the feature name, and the top 3 principal components similarly extracted (“STAR-PC”).

Given the gene expression and covariate matrices, features that explain a significant proportion of expression variance in a non-trivial subset of genes were extracted using a forward-backward regression approach (see supplemental methods). This approach identified the features “seq_pc1”, “seq_pc2”, “seq_pc3”, “SMRIN”, “SMEXNCRT”, “Number_of_splices_GT/AG”, “TRISCHD” and “DTHCODD” as significant features, with no significant interactions between these features or between any of these covariates and tissue type.

Because there were no significant cross-terms between tissue and covariate, all tissues were combined for the removal of covariate effects. A linear model (expr ∼ tissue + covariates - 1) was applied, with a separate intercept (mean) for each tissue. The covariate effects were removed, while the estimates of mean expression per tissue were retained.

## Forward-backward covariate selection using MARS (earth)

A key step in the treatment of RNA-seq data is identifying what technical or biological covariates are strong drivers of measured expression. RNASeqQC produces a large set of alignment metrics derived from the aligned RNA-seq bams. We combined these with the splicing metrics output by STAR. Separately, each of these data were scaled and the top 5 PCs calculated to summarize the bulk of the technical covariate distribution, producing an additional 10 potential covariates. This final set of technical covariates are combined with the sample-level individual-level information provided by GTEx (ischemic time, age, biological sex, RIN, ethnicity, race).

We then used the ‘earth’ package in R to select covariates that explained a large amount of expression variance across many genes. We set the parameters so that no non-linear splines were used, but that cross terms up to degree 3 were allowed, enabling the model to select tissue-by-covariate or covariate-by-covariate effects.

earth builds a forward model by selecting the covariate (or cross term) which most improves the total R^2 across all genes considered; and when a diminishing-returns threshold is reached (for us, an improvement of 0.01), prunes the terms using a penalized R^2 heuristic.

We ran earth 100 times on a random sample of 1,000 genes; each run producing an estimate of variance explained for all covariates (covariates *not* included in the model are assumed to explain 0% of expression variance). We summarized the impact of each covariate by taking the upper 20% of the variance explained (**figure S1a**). Any covariate whose summary estimate was >5% variance explained was included in our final model for covariate correction. For group variables (such as tissue); if any subgroup exceeded the variance explained threshold, then the entire group variable was selected.

Notably, no cross-terms exceeded the threshold for variance explained, suggesting that we could perform covariate correction simultaneously across all tissues. The lack of region-by-covariate effects may be due to the fact that the library preparation batches and sequencing batches are well-balanced across brain regions.

## Tissue hierarchy

The median expression of all genes across a given tissue is taken as the *exemplar* of said tissue. These exemplars (12 in all) are hierarchically clustered into the tissue hierarchy observed in figure 1 using Euclidean distance and single-linkage hierarchical clustering.

## Module construction

### Robust WGCNA

Robust rWGCNA^clxxii^ was applied to each brain tissue independently. Briefly, the power parameter is selected as the smallest power (between 6 and 20) which achieves a truncated r^2 of >0.8 and a negative slope. Then, 50 signed co-expression networks are generated on 50 independent bootstraps of the samples; each co-expression network uses the same estimated power parameter. These 50 topological overlap matrices are then combined edge-wise by taking the median of each edge across all bootstraps.

The topological overlap matrices are then clustered hierarchically using average linkage hierarchical clustering (using ‘1 – TOM’ as a dis-similarity measure). The boostraps are used to determine cut height as follows: multiple cut-heights are considered (0.9 to 0.999, by 0.005); and for each cut the within-module correlation of TOMs is considered. For the top 8 modules by size (fewer if fewer modules are produced), the consensus and each bootstrap TOM is subset to the genes within each module, and the correlation between bootstrap and consensus is computed. The median (within module, across bootstraps) of these consensuses is computed, and the mean of these summaries is taken to be a measure of ‘goodness’ for the cut. The cut height which maximizes this metric is taken to define the initial modules.

These initial modules are then merged via ‘mergeCloseModules’ in WGCNA, which hierarchically re-clusters modules based on the module eigengenes, using the correlation-based adjacency as a dis-similarity matrix. Modules with a distance of < 0.35 are merged together into a combined module.

### Aggregating co-expression

At each merge of the hierarchy, a single round of consensus topological overlap is performed. Each pair of genes has two descendent edges, and the parent edge is estimated as the 80^th^ percentile between the two (i.e. for x < y; p = 0.2 x + 0.8 y)). This process proceeds up the tissue hierarchy until a single network TOM remains.

### Consensus labeling

After construction of co-expression networks from all tissues and splits, modules have been defined for a total of 21 groups (BRNACC-BRNSNA, BROD, CTX, CBL, BGA, STR, NS-SCTX, SCTX, NCBL, WHOLE-BRAIN), yielding over 300 overlapping modules. The overlapping nature of these modules motivates labeling each module in terms of a hierarchy group, allowing one to identify (say) BRNHYP-M2 and BRNCTX-M7 with the module group WHOLE-BRAIN-M3.

To perform this labeling, similarity matrices are computed. First, the module eigengenes for all modules (regardless of origin) are computed within every tissue, and the correlation matrix (using ‘bicor’) is computed for each module for each tissue. This produces an (all modules) x (all modules) matrix for each tissue. The consensus eigengene similarity (“E”) between two modules is chosen as the component-wise maximum of all of these matrices. The second similarity matrix is the standard Jaccard similarity (“J”) between module gene lists. These similarities are combined into a dis-similarity matrix D = 1 - (E + 3*J)/4, which is used to hierarchically cluster (average linkage) these modules.

Module groups are defined by cutting the dendrogram at a height of 0.35. This process results in a set of module clusters, each of which has a “level” in the brain tissue hierarchy (for instance, a cluster of BRNCTXBA9-M4, BRNCTXB24-M2, CTX-M7 would have the level “CTX” as the top-level of the tree represented is CTX). The “representative” of the module group is taken to be the module at the highest (most rootward) level of the tree – and if there are two, the larger of the two. A second round of clustering is performed by removing all modules in the group (except for its representative) from the dissimilarity matrix, and re-clustering only the group representatives. This process repeats until there are no additional merges. Finally, each module is labeled with its group representative; for instance “BRNCTXBA9-M4” would receive the label “CTX-M7”, because it shares its highest similarity with the consensus cortex module M7.

In addition, we re-named and abbreviated modules: “BW” for brain-wide, “NCBL” for non-cerebellar, “NS.SCTX” for non-striatal subcortex, “CEREB” for Cerebellum; and the GTEx tissue names were abbreviated to clear region codes: ACC, AMY, B24, BA9, CBH, CBL, CDT, HIP, HYP, PFC, PUT, SNA.

## Preservation

We consider two module preservation statistics: the classical Z-summary^clxxiii^ and a leave-one-gene-out neighbor statistic. For the classical Z-summary; module statistics such as the mean gene-gene correlation in the module, the correlation-of-correlations across datasets, the variance explained by the first module PC, and other metrics are computed for each module (in both the original and comparison dataset); and compared to 100 random (via permutation) modules of identical size. Each observed statistic is converted to a Z-score, and these are averaged to generate a final summary, for which large Z-scores are indicative of replication of the underlying biological signal.

The neighbor statistic (“Z-AUPR”) is strongly influenced by the single-cell statistic MetaNeighbor^clxxiv^. Briefly, a *k-*nearest-neighbor network is built in the comparison dataset (we use k=15), and we impose the module labels from the reference dataset. For each gene, we compute the proportion of its neighbors (again, in the comparison dataset) whose labels match its own. Note that if this proportion is > 0.5, then this gene *would* be assigned the same label in the comparison dataset as the reference dataset under a neighbor-voting scheme. Using these scores, we can compute an AUPR for each module. We repeat this approach for 100 permuted modules (and, unlike the WGCNA permutation, we split genes into connectivity deciles, and permute only within decile), and use this baseline to convert observed AUPR to Z-scores. As with the classical Z-summary, high Z-AUPR is indicative of replication of underlying biological signal.

## Module comparisons

We considered three alternatives to WGCNA for network building and module identification: ARACNe, GLASSO, and von-Mises-Fisher clustering.

ARACNe was run with default settings (10 permutations, FDR of 0.05); and genes filtered by ARACNe (for having no significant edges) were placed into a background ‘grey’ module. The resulting network was imported into iGraph^clxxv^ and modules identified by Louvain clustering.

As sparse inverse-covariance estimation is computationally intensive, we took an approximate approach. First, we partitioned the genes into initial groups of approximate size 1000 using k-medioids clustering. GLASSO was applied independently to each group to estimate a blockwise precision matrix. Within each block, the penalty parameter was selected using StARS^clxxvi^, targeting an edge instability of between 0.05 and 0.1. Genes with no partial correlation to any others were grouped into a background ‘grey’ module. The remaining network was imported into iGraph and modules identified by Louvain clustering.

vMF clustering, unlike the other approaches, does not build a network, but seeks to identify gene clusters directly. Gene expression vectors were pre-processed by transforming their values into ranks (across samples) and normalizing them to unit norm. In this way, an inner product between two gene vectors is effectively their Spearman correlation. The resulting data is modeled as a collection of draws from an *n*-dimensional mixture of *k* von-Mises-Fisher distributions (where *n* is the number of samples). The model was fit using the R package movMF^clxxvii^ for *k* varying from 8 to 50. The final choice of *k* came from the model that maximized likelihood – 2 * ndim * k; and module assignments were determined from the most likely mixture probability (or ‘grey’ if that probability was less than 0.8).

## Whole-brain module comparisons

Beyond comparing modules within each tissue, we sought to compare our hierarchical WGCNA modules with an orthogonal approach for building consensus modules. As consensus modules built from methods already similar to WGCNA would certainly produce similar consensus modules, we considered an alternate approach: tensor decomposition.

First, we built a fully imputed (gene x brain x region) tensor by using probabilistic PCA to impute missing samples within every (brain x region) submatrix for each gene. We then applied CANDECOMP to this tensor to produce 150 feature triplets: {(gene x 1), (brain x 1), (region x 1)}. We treated the gene-level features as a (gene x 150) feature matrix, and ran t-SNE to embed the genes in a 2-dimensional space.

While this embedding did not show distinct visual clusters, it clearly showed regions of high and low density, likely corresponding to modules. Given this intuition, we applied the DBSCAN clustering algorithm, producing a set of 30 whole-brain modules.

We found that the ribosomal, glial, and choroid-plexus modules were in one-to-one correspondence with TD-DBSCAN modules (figure S1d, e), and that the neuronal WGCNA modules correspond to multiple TD-DBSCAN modules, with statistically significant overlaps. Visually, the WGCNA modules are localized in the embedded tensor-decomposed space, strongly suggesting that the modules are not driven by the specifics of WGCNA, nor are they induced by the structure of hierarchical merging; but rather that these genes are grouped together by disparate approaches because of an underlying biological signal.

## Learning curves

To examine how module identification and specificity changes as a function of the number of samples, we combined samples from similar tissues to increase the maximum N: we combined the cerebellar samples into one larger group (N=122), and we also grouped the cortical samples (PFC, B24, BA9) together with hippocampal samples into a second group (N=304).

“Reference” modules for these groups were determined by applying rWGCNA to the full dataset. We down-sampled the group to a smaller set of samples of size n = 25, 50, …, N and performed rWGCNA on the smaller set. We repeated this process 10 times, generating 10 networks and module assignments for each sub-sampling of the full dataset.

Because two clusterings should be considered identical up to renaming the labels in one or the other datasets, we use module co-clustering as a measure for accuracy, precision, and recall. Within the reference (whole group) dataset, we extract the top ‘hub’ gene from each of the modules, and the list of genes co-clustered with that hub gene (i.e. the other members of its module). For a given reference module, within a sub-sampled dataset, we have

Recall = (# ref hub co-clustered genes also co-clustered in subsample)/(# ref hub co-clustered genes)

Precision = (# ref hub co-clustered genes also co-clustered in subsample)/(# subsample co-clustered genes)

In effect, these are precision/recall statistics for the hub gene co-clustering indicators. If two reference modules fail to separate in a sub-sample (a typical failure mode), the result is slightly higher recall, but far worse precision.

## Single-cell data

Quantified single-cell data was downloaded from http://mousebrain.org^clxxviii^ (mouse) and subset to only cells from the CNS (without spinal chord); and GEO GSE97942^clxxix^ was downloaded for human. These data were log-transformed log(1 + x) for counts and log(0.005 + x) for TPM; and the cell type labels from the respective publications were used for all subtype analyses. Absolute expression values were taken as the mean expression of a cluster; and relative expression was obtained via

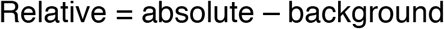

Where the background expression is the average expression of a gene over all cells. To incorporate gene variance information into relative expression, the *relative expression rank* is defined as the lower end of a small confidence-interval for the difference in means:

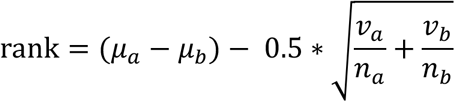

kME enrichments are based on the correlation between module kME and the relative expression rank within a given cell type.

## Cell-type enrichment and single-cell data

For kME-based enrichments (such as those in figure 2), the shaded region of the figure represents the standard error around the estimated functional relationship between kME and relative expression rank. In all cases it is visually apparent that these lines deviate from 0 by a factor far exceeding 2.5 times their standard error (p ∼ 0.006).

For gene-set based enrichments such those presented in the text, and those in figure 3, cell type markers were obtained from several sources^clxxx,clxxxi,clxxxii,clxxxiii,clxxxiv,clxxxv,clxxxvi,clxxxvii^ representing various studies performed both in mouse and in human. The statistical test is a logistic regression using the model

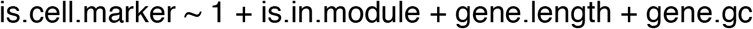

adjusting for gene length and GC. We test that the coefficient for module presence is significantly different and greater than zero, implying an enrichment (as opposed to depletion) of cell-type related genes.

This test is performed independently on cell type markers from the various studies, and FDR adjusted across all tests.

## Defining mouse orthologs to human genes

The ensembl API was used, through biomaRt, to query human genes with associated mouse orthologs and the type of orthology; and visa versa. These queries enabled defining genes as one-to-one orthologs, one-to-many orthologs, many-to-many orthologs, or non-orthologous. The ensembl API was also used to obtain human-mouse dN and dS values; and the ratio dN/dS calculated, with 0/0 treated as 0.

## Module Imputation

For our lncRNA analysis, we imputed whole-brain modules into an independent RNA-seq dataset^clxxxviii^ by i) splitting the data into BA9 and BA41-42-22 regions, ii) Calculating module kMEs within each region, and iii) Averaging across the two regions. This generates a set of 11 features (average within-region kME to each module) for each gene. The overlapping genes between the GTEx modules and control brain expression were used as labels to fit a boosted trees classifier (using the R package xgboost with 2000 trees and a learning rate of 0.025). Non-overlapping genes (which contain most lncRNA and a set of held-out, matched protein-coding genes) are assigned to modules via the prediction of the fitted classifier. Using cross-validation on the matched protein-coding genes, we estimate that the sensitivity and specificity of this approach are 0.63 and 0.53 for BW-M6, with sensitivity ranging from 0.25-0.7 and specificity from 0.2-0.8 across other modules. The most common misclassification (>60%) results from assigning a ‘grey’ gene as in the module, or a BW-M6 gene as ‘grey’.

## Human-specific modules

To define modules exhibiting human-specific differential expression, we obtained the modules and human-specific differentially-expressed gene list from Sousa2017.^clxxxix^ We subset only to modules flagged as showing inter-species heterogeneity, and computed enrichment p-values and FDR values by Fisher’s exact test, using the intersection of all GTEx-ascertained genes and Sousa2017-ascertained genes. This resulted in a set of 25 modules with enrichment FDR<0.1 for human-specific differentially expressed genes.

Using the same statistical approach and background gene set, we then tested for significant overlaps between the 311 GTEx modules and the 25 human-specific modules, identifying 10 with an enrichment FDR < 0.1. These overlaps are plotted in figure 2(g). Not every module in brain-wide module sets necessarily overlapped at FDR<0.1; so the figure reflects the proportion of modules within brain-wide module sets that show such an overlap. Furthermore, because hypothalamus and substantia nigra were not profiled in Sousa2017, these regions (and the NS.SCTX region) were excluded from this fraction calculation (but not from the initial overlap tests and FDR correction).

Sousa2017 also lists cell types in which these modules are expressed. These are summarized in figure 2(g). Expression is listed for cell types Ex1-Ex8 and In1-In8; for space this is collapsed to the fraction of Ex and In in which the module is expressed, so a gene expressed in In4 and In2 would receive a value of 0.25 for the “In” group.

## GO enrichment

Gene ontology enrichment is performed competitively, with covariate correction, using logistic regression. Briefly, each GO category is treated as a binary variable (1 for genes in the category, 0 for genes not in the category – only genes ascertained in our gene expression matrix are part for the regression). Modules are also treated as binary. We include as covariates the average gene expression across all tissues in the brain, the gene GC content, the log gene length, and the gene expression reproducibility (see below). The GO enrichment model is then

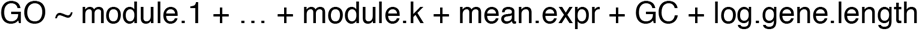

And is fit using logistic regression. If we detect that convergence fails, an L2-regularized logistic regression is instead applied (using ‘brglm’). The enrichment p-values are taken to be the statistics that reject (β_i_ ≤ 0) for all β_i_ corresponding to a module indicator.

The enrichment p-values are adjusted for all ontologies.

## Meta-GSEA

To aggregate enrichment results (such as GO) from the module level to the module set level, the GO p-values are treated as independent p-values, and Fisher’s method is applied: For a given ontology category, a χ^2^ value is calculated as -2 * log(p_1_*p_2_*…*p_k_), where the product is taken across modules in the set. In the case of independence, this statistic has 2\**k* degrees of freedom; allowing a p-value to be calculated. Because the modules in a set overlap by construction, the resulting statistics are not calibrated probabilities, and are referred to as “scores” or “rankings,” and should not be interpreted as reflecting significance. In nearly all cases, the highly-ranked consensus ontology had been significant in one or more of the modules within the set.

The meta-GSEA applied to generate supplemental figures 5b,c was to identify the genes within the regional BW-M4 modules (e.g. PFC-BW-M4) with MAGMA Z-scores > 3.0 (SCZ) or 2.5 (ASD). This generated an indicator variable which was then used to perform gene ontology, using the BW-M4 genes as a background; generating p-values for each ontology. Meta-GSEA was applied to these p-values, generating a score for each ontology, plotted in 5b,c.

## pLI enrichment

Gene pLI scores were downloaded from the ExAC consortium release^cxc^, and a gene was considered likely to be LoF-intolerant if its pLI score was 0.9 or higher. Enrichment for “hard” module membership (i.e. comparing two gene lists) is performed via Fisher’s exact test on the contingency table between module membership and LoF-tolerance/intolerance. “Soft” module enrichment (i.e. based on kME) is computed via a Brownian Bridge statistic.

The genes are ranked by their module membership (kME); and the proportion of all genes which are likely LoF-intolerant (the pLI rate, *r=P/M*) is computed. At a given quantile *q* of genes, we tabulate how many of the first *q * M* genes are LoF-intolerant; and denote this cumulative sum by *Cs(q)*. The expected number of LoF-intolerant genes is Ne(q) = *q * P = q * r * M.* For large M, this cumulative sum converges to a scaled Brownian motion with drift *r;* and has variance V(q) = *q * (1 - q) * M * r * (1 - r)*. Z-scores for this cumulative sum at each *q* are given by Z(q) = (Cs(q) - Ne(q))/√V(q). An excess of LoF-intolerant genes occurs when *min_q Φ*(*Z*(*q*)) < 0.05. For clearer visualization, we plot (Cs(q) - Ne(q)) and 2.17 * √V(q) as functions of q.

We also used a generalized additive models (“GAM”) and a generalized linear models (“GLM”) to verify findings of constraint. In these cases we applied the (logistic) model

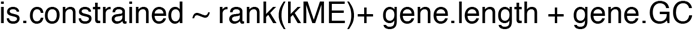

and found that, for the whole-brain modules, these enrichments were so strong that the three methods were in 100% concordance. The results of the linear models did not change substantively when using competitive as opposed to marginal enrichments.

For supplemental figure 4 (enrichment in pLI and o/e bins), the odds ratio and p-values were computed using a Fisher Exact Test between module membership, and bin membership.

## PPI enrichment

We use the InWeb PPI database^cxci^ (brain tissue) for a source of defined protein-protein interactions, with a confidence threshold of 0.2 used as a cutoff for a particular interaction. PPI prediction is treated as edge-related data, where the response variable is binary (presence/absence of PPI), and the predictors the following collection of data relevant to that edge: the (PPI) connectivity of its first vertex, the (PPI) connectivity of its second vertex, the product of kMEs of its vertices (for each module), the product of the GCs of its vertices, and the product of the reproducibilities of its vertices. Or:

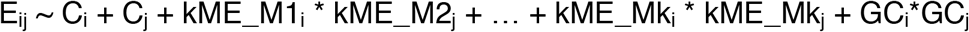

This equation encodes the model that gene pairs which are mutually close to a given module are more likely to physically interact. The logistic model is fit using ‘statsmodels’ in python, and the hypotheses βi ≤ 0 is assessed for each βi corresponding to a module.

## Regional contrast test

The Regional Contrast Test is a multivariate test of significance for

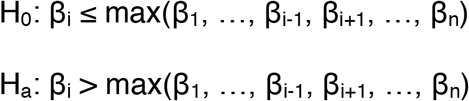

This statistic corresponds to a multidimensional integral, with infinite limits on all coefficients other than β_i_, and taking max(β_1_, …, β_i-1_, β_i+1_, …, β_n_) < β_i_ < ∞. Because of the large numbers of degrees of freedom in this regression, we treat the variance-covariance matrix (Σ_β_^(ML)^) of the β vector as giving the true sampling covariance of these parameters, and perform Monte-Carlo integration by drawing 50,000,000 samples from the multivariate normal distribution N(β, Σ_β_^(ML)^) using the R package *fastmvn*.

The above statistic works for testing each tissue against all others. A grouped version of the test is a simple extension, which considers several β in tandem. For simplicity we assume the indexes for the group are the first *k* coefficients, then the comparison becomes:

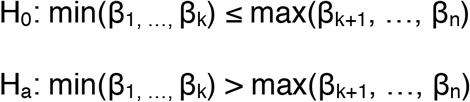

This only changes the integration limits to (for j ≤ k) to max(β_k+1_, …, β_n_) < β_j_ < ∞; and we use the same Monte-Carlo approach as before.

## Post-hoc tests for module enrichment use Fisher’s exact test on the contingency table

|  | Significant<br>(RCT) | Not significant<br>(RCT) |
| --- | --- | --- |
| In module | A | B |
| Not in module | C | D |

## Isoform specificity from sorted cell data

RNA-sequencing data was obtained from GSE73721 (SRA project SRP064454) and quantified at the isoform level with Kallisto (mouse gencode release M16). These data included sorted populations of astrocytes, oligodendrocytes, endothelial cells, a single neuronal population, and a whole-tissue background. Relative isoform expression were obtained as described in “Single-cell data,” with the background set to be the average expression across the whole-tissue background samples.

## Isoform switching and validation

Isoform-level TPM values (produced by RSEM) were corrected using a linear model with the same covariates used for correcting gene expression TPMs. Subsequently, each isoform expression (within tissue) was correlated to brain-wide module eigengenes computed within the tissue, and the mean correlation across tissues taken as an estimate of module membership for the isoform.

To determine an appropriate kME threshold, we evaluated the impact of thresholding on cell type enrichments. Each threshold produces a set of isoforms within a module; and each isoform can be annotated with the cell type marker status of its parent gene. Fisher’s Exact Test produces an odds ratio and p-value for cell-type enrichment at each threshold. We found that a threshold of 0.45 produced a 15-fold enrichment for both astrocyte and oligodendrocyte markers when looking at kME to their respective modules (M6 and M7); but that when increasing this threshold the odds ratio for oligodendrocytes did not substantially change, while the astrocyte odds ratio increased **(figure S7)**. Based on this we defined the threshold for isoform module membership at 0.45 kME. In the case where an isoform has >0.45 kME to multiple modules, module with highest kME is selected.

An “isoform switch” is defined as two sister isoforms having membership to different modules.

## Western Blot Analysis

Human iPS cells were differentiated into cortical glutamatergic-pattern neurons (GPiN) according to Nehme 2018,^cxcii^ and samples extracted at days 0, 16, 21, and 31. Human astrocytes were used as an outgroup. IP was performed using an ANK2-specific monoclonal antibody S105-17.

## De-novo variant enrichment

Denovo-DB^cxciii^ was used to extract lists of genes harboring *de novo* variation linked to ASD and Schizophrenia. The v1.5 of the database was obtained on 02-17-2018, and we filter for “PrimaryPhenotype=autism” (or, separately, “PrimaryPhenotype=schizophrenia”) and “FunctionClass” as one of “frameshift”, “frameshift-near-splice”, “splice-acceptor”, “splice-donor”, “start-lost”, “stop-gained”, “stop-gained-near-splice”, or “stop-lost.”

Module enrichments are calculated via Fisher’s Exact Test, using the contingency table formed by cross-tabulating module presence/absence with presence/absence on the denovo-db gene list.

As the denovo-db is a broad collection of *de novo* mutations in affected individuals and does not curate these variant lists on the basis of total evidence, we consider two additional data sources for alternative enrichment scores. First, we consider the curated list of SFARI genes of rank S, 1, 2, or 3; and perform enrichment on the resulting likst.

## Second, recent work from our lab^cxciv^ computes transmission and de-novo association

Bayes Factors for 18,472 genes. We regress the log Bayes Factor against module presence/absence and look for a significant, positive coefficient.

## GWAS variant enrichment

Enrichment for GWAS signal was performed through the use of MAGMA^cxcv^ gene set analysis. Briefly, variants were mapped to genes on the basis of genomic distance, while taking chromatin contact maps from adult brain Hi-C^cxcvi^ into account. MAGMA was used to generate gene scores and LD-based covariances. Subsequently, MAGMA’s gene set analysis was used to compare the distribution of gene scores between modules and the background set of ‘grey’ genes.

8 GWAS studies were considered in this analysis: The iPsych and PGC cross-disorder GWAS studies (accounting for ASD, SCZ, and cross-disorder), Alzheimer’s disease, multiple sclerosis, and educational attainment.^cxcvii,cxcviii,cxcix,cc,cci^

## Differential preservation analysis

Modules defined in the GTEx tissue samples were assessed for their preservation in ASD case samples and (separately) in normal samples. These produced a pair of preservation Z-scores per module. We defined differential preservation to be cases where the control Z-score is preserved (>3) while the ASD Z-score is not preserved (<3).

## Core/periphery enrichment within networks

### Simulation of network genetic architecture

#### Simulation

10,000 causal variants are simulated with frequency parameters estimated from human populations,^ccii^ and distances drawn from a binned Beta distribution:

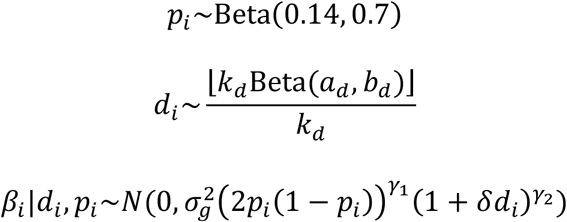

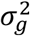 is arbitrary and set to 1; *k_d_* is arbitrary so long as it is greater than about 5, and is set to *k_d_*=12; *a_d_*, *b_d_*, *γ*_1_, *γ*_2_, and *δ* are model parameters. Recent results from the UK Biobank suggest that a value of *γ*_1_ = −0.4 is reasonable for a polygenic trait (height=- 0.45, education=-0.32, blood pressure = -0.39) and is fixed to this value. Architectures were simulated on a grid of *a_d_*, *b_d_*=1,1.5,…6; *δ*=1,1.2,…,2.6; *γ*_2_=-15,-10,-7,-5,-2. Notably for any values of *a_d_*, *b_d_*, *δ* and *γ*_2_ can be found such that D_1_ explains >40% of the heritability. <u>Errors-in-distance</u>: Here the above simulation of distance is replaced by a normal copula (where 20% error corresponds to r=0.8 – this is a purposeful under-estimate, as r^2^=0.64 so the latent error is more like 36%):

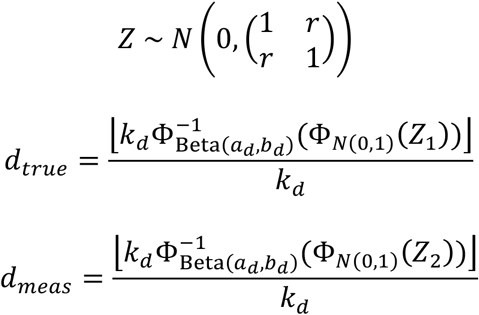

When simulated from a network, first a set of K=1, …, 10 hub genes are simulated with the constraint that no pair can be directly connected by an edge. These form initial communities of size 1. For the remaining 40 core genes, a community is selected at random, a community member is selected at random, and a neighbor is selected at random and added to the community and to the set of core genes. These form the basis of *d_true_*, which is taken as the minimal path distance to any core gene. For *d_meas_* the communities are distorted by removing M=1,..,10 core genes at random; or by adding K=5,10,…,25 non-core genes at random.

#### Normalized effect sizes

Identifying the effect size of an empowered 5% frequency GWAS variant happens through three steps: (i) Estimating the liability distribution; (ii) Mapping case/control frequency differences to effect sizes (iii) Estimating power.

i. Liability Distribution: A 5000×10,000 genotype matrix X is sampled independently, with frequencies given by the previously-simulated vector f, and 5,000 genetic liabilities are generated by lg=Xβ. These liabilities are used to estimate parameters for a T-distribution using ‘fitdist’ from the R package ‘MASS’; the degrees of freedom are reduced by 25% to account partially for rare variants not sampled in this population of 5,000; and these parameters used to generate 400,000 genetic liability scores. These are converted to total liability scores by adding noise l = lg + N(0, σe); with σe chosen so that the heritability is 0.85.
ii. Frequency-ratio-to-effect: The goal is to estimate the ratio paff/punaff for a variant with a frequency pi and effect βi. The genetic liabilities lnew = l+xβi with x ∼ binomial(2, pi) are computed for 400,000 simulated individuals. As 10,000 variants contribute to l, the addition of xβi is assumed to have a minimal effect on heritability. Case/control labels are defined by lnew ≥ quantile(lnew, 0.95) so that the disease prevalence is 5%, and the empirical frequency mean(xaff)/mean(xunaff) is taken as an estimate of the ratio paff/punaff. Fixing pi=0.05 and varying βi produces an empirical and invertible map from variant effect to frequency ratio.
iii. Estimating power: Given an effect size βi, the case and control frequencies for a p = 0.05 variant are obtained from (ii). 5000 case and 5000 control genotypes are sampled according to the corresponding frequencies, and a two-sided T-test performed by ‘t.test’ in R. 1,000 simulations are performed, and the number of times the T-test p-value achieved a Bonferroni-corrected p-value of 0.1/10,000 (the number of causal variants) was tabulated.

### Network construction and computation of d(G)

#### Co-expression Networks

Within co-expression networks, the raw co-expression (cosine) distance is used to define gene-gene distances. In addition, a sparse ε=2.5%+1-NN graph is calculated as follows: the cosine distance graph is subset to only the 2.5% smallest edges, and any singleton genes are connected to their closest neighbor. This graph is treated as unweighted, and not necessarily connected. Cross-component distances are treated as 1 + the maximum observed within-component distance. This is referred to as “sparse distance.”

Module hub genes are defined as the 2.5% of module genes with largest kWithin values (minimum 5). Distances between a gene and a module is computed as (i) 1 – kME; (ii) mean cosine distance to a module hub; (iii) minimum cosine distance to a module hub; (iv) mean sparse distance to a module hub; (v) minimum sparse distance to a module hub. When using arbitrary gene sets as core genes, (ii)-(iv) are be computed with respect to the gene set in place of module hubs.

#### Transcription Factor Binding Networks

Bipartite transcription factor binding graphs were obtained from regulatorycircuits.org, and converted to a similarity network as in Marbach2016. Briefly, the probability weights are taken as edge weights, and the random-walk kernel *K=*(*I+W*)^4^ with *W* the symmetrically-normalized Laplacian *D^-1/2^AD^-1/2^* of the adjacency matrix; and converted to a dissimilarity via 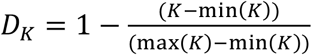. A natural set of “core” genes on this network are the most highly-connected genes of *K*; of which the top 25 are taken. Distances are either the mean or minimum path distance under D*_K_*.

#### Protein-protein interaction networks

InWeb^cciii^ was used for the protein-protein interaction network. The refined brain-PPI network was obtained from the resource, and a confidence of 0.05 required for an edge to be defined; and the interactions were converted into a binary matrix. Distances were defined as either the minimum or mean path distance in this network. As with TFBNs, the natural set of hub genes are the most connected genes, of which the top 25 are taken.

### Hub genes and empirical core genes

Core gene sets which define distance (“proposal set”) are taken to be either collections of network hub genes, or the top 10 or 20 genes (by Bayes factor) from each of the three studies (separately). The core gene sets which define the statistic Φ (“evaluation set”) are taken to be the top 25, 35, 50, 75, or 100 genes from each of the three studies. To restrict attention to directly causal (e.g. non-regulatory) genes, as the omnigenic model suggests, the core genes are also filtered to remove known transcription factors,^cciv^ DNA-binding proteins, RNA-binding proteins, and non-coding RNA. Without this filtering, values of Φ still fall below 50% for brain co-expression networks, but achieve 70% for blood co-expression. Φ statistics are calculated for only for evaluation sets where, after excluding those genes also in the proposal set, noncoding genes, DNA-binding proteins, known transcription factors, and RNA binding proteins, at least 15 genes remain.

### Significance calculation for Φ

Because Φ reflects a partitioning of a subset of genes, a significance value can be calculated by Fisher’s Exact Test. As a specific example: the overlap of genes between two studies is 15902 genes. After computing network distances, the top decile contains 1590 genes. Imagine that the core gene set (after excluding non-coding, non-regulatory genes) contains 31 genes, and 12 of these overlap the set of 1590. The contingency table 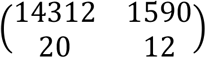 reflects this observation, and has a p-value of 0.001.

---

**Supplemental Figure 1: Related to figure 1.**
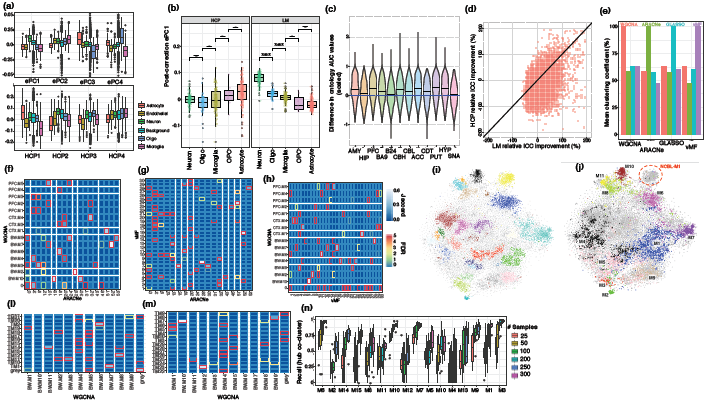
(a) ePC and HCP loadings onto canonical cell type genes, showing significant heterogeneity of loadings across cell types. (b) ePC loadings after covariate correction using HCP and LM base correction, showing that cell type heterogeneity of the 1st component of expression is lost after HCP correction. (c) Network-based GO prediction accuracy for each brain region. The same gene holdouts are used in 10-fold cross validation, generating 10 values for the AUC difference of each GO category, which are used to generate a Z-score for the expected AUC difference. (d) Relative improvement to the integrated correlation coefficient (Parmigiani 2012; suppl methods) for BRNHYP genes, for linear model and HCP based corrections. (e) Pairwise co-clustering statistics for the 4 algorithms compared in figure 1. X-axis denotes which modules are taken as the reference set. (f-h) Pairwise module overlaps between 3 of the 4 algorithms compared in figure 1 (GLASSO yielded too many modules to visualize here). (i) t-SNE embedding of gene features from whole-brain tensor decomposition, colored by DBSCAN clusters. (j) As (i), but colored and annotated with whole-brain modules. (l,m) Overlap between whole-brain consensus and tensor-decomposition+DBSCAN modules. Color scheme as in (f-h). (n) Within-module recall curves for hub-gene co-clustering, demonstrating that at 100 samples, the recall is above 50% for most modules.

**Supplemental Figure 2: Related to figure 2.**
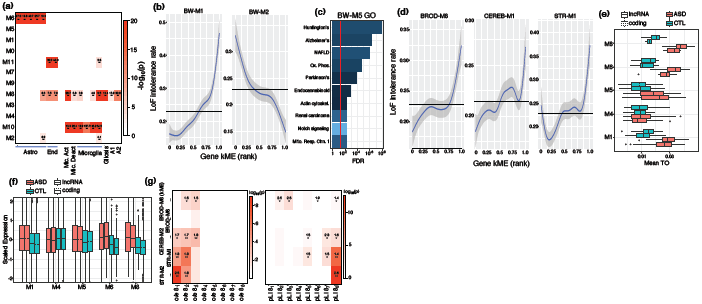
(a) Cell-type marker enrichment for brain-wide modules, extended with markers of microglial activation and deactivation, and markers of reactive gliosis and A1/A2 reactive astrocytes. (b) Plots of the marginal rate of LoF-intolerant (pLI>0.9) genes, as a function of BW-M1 (most enriched) and BW-M2 (most depleted) kME. (c) Gene ontology enrichment for BW-M5. (d) Marginal LoF-intolerance rates, by gene kME, for neuronal subtype modules. (e,f) Module mean topological overlap, and gene expression, for 5 whole-brain modules in ASD cases and matched controls (Parikshak 2016). The case/control difference in lncRNA is closely matched by the same difference in randomly-selected, matched coding genes. (g) LoF-intolerance enrichment for neuronal subtype modules, using pLI and o/e bins as response variables, and a linear model correcting for gene GC and length. All modules except BROD-M8 show strong enrichment, and BROD-M8 shows enrichment when using soft-membership instead of hard membership.

**Supplemental Figure 3: Related to figure 3.**
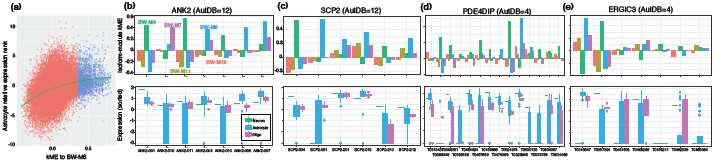
(a) Replicate of main figure 3(b) in astrocytes, showing a strong positive relationship between astrocyte module membership, and relative expression in astrocyte cells. (b-e) Relationship between module kME and cell type relative expression for transcripts across 4 neuron/astrocyte isoform switch genes, demonstrating concordance between high kME, and high relative expression.

**Supplemental Figure 4:**
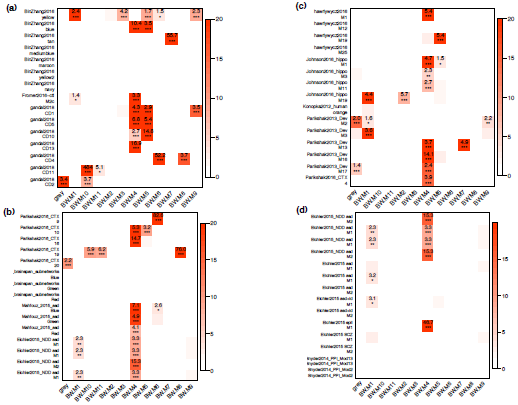
Published module overlaps. (a-d) Overlaps between published modules and the consensus whole-brain co-expression modules identified in this paper, demonstrating that the majority of modules show a high overlap, particularly to the neuronal module BW-M4. These modules were been selected because of published enrichment for neuropsychiatric disease risk genes. (supplemental methods).

**Supplemental Figure 5: Related to figure 5.**
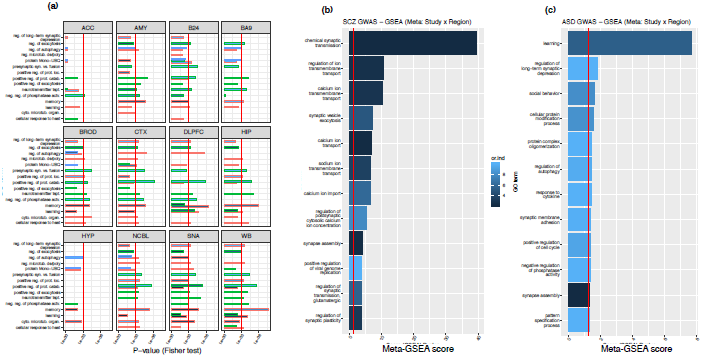
(a) Gene ontology enrichments for module set BW-M4 across all regions in which a BW-M4 module is present. (b) Meta-GSEA scores for significant MAGMA genes in BW-M4 across all tissues, implicating synaptic transmission and calcium transport as neuronal dysfunctions in SCZ.

**Supplemental Figure 6:**
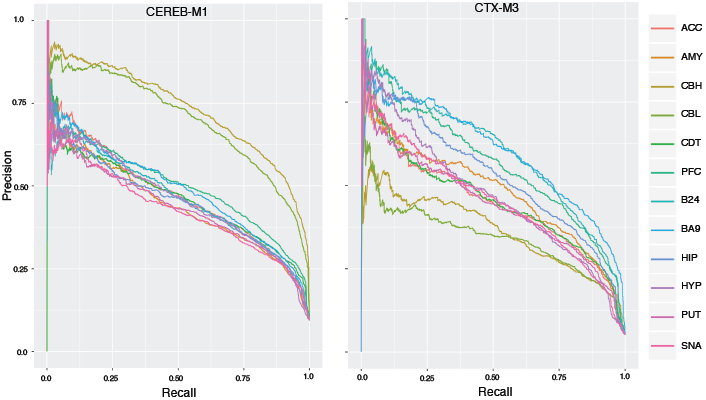
Regional AUPR curves for CEREB-M1 and CTX-M1. *Left*. Nearest-neighbor precision-recall curves for CEREB-M1 labels across all region-level co-expression networks; showing significantly higher AUPR for cerebellar regions, but substantial AUPR for all remaining regions. *Right*. Nearest-neighbor precision-recall curves for CTX-M3.

**Supplemental Figure 7:**
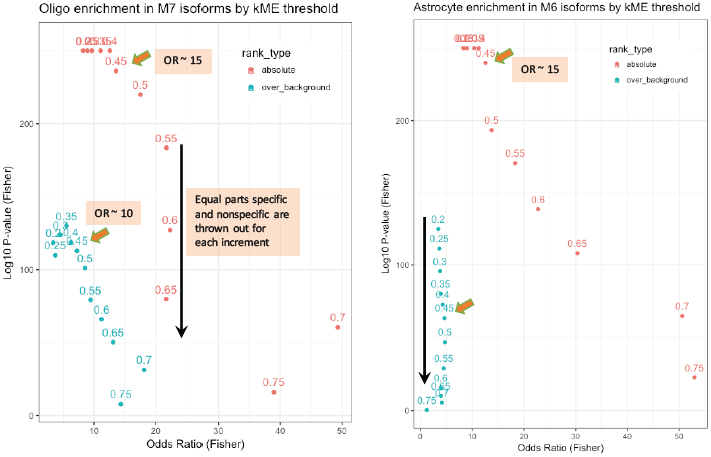
isoform module assignment threshold, related to figure 3. Fisher’s exact test of the contingency of “assigned to module” and “top-ranked cell type marker” for varying kME thresholds for (*left*) oligodendrocytes and (*right*) astrocytes; for marker rankings based on both absolute and relative expression within the cell-sorted data. Thresholds in the range 0.45-0.55 appear to balance significance and odds ratio across absolute and relative rankings.

**Supplemental Figure 8: Related to figure 7.**
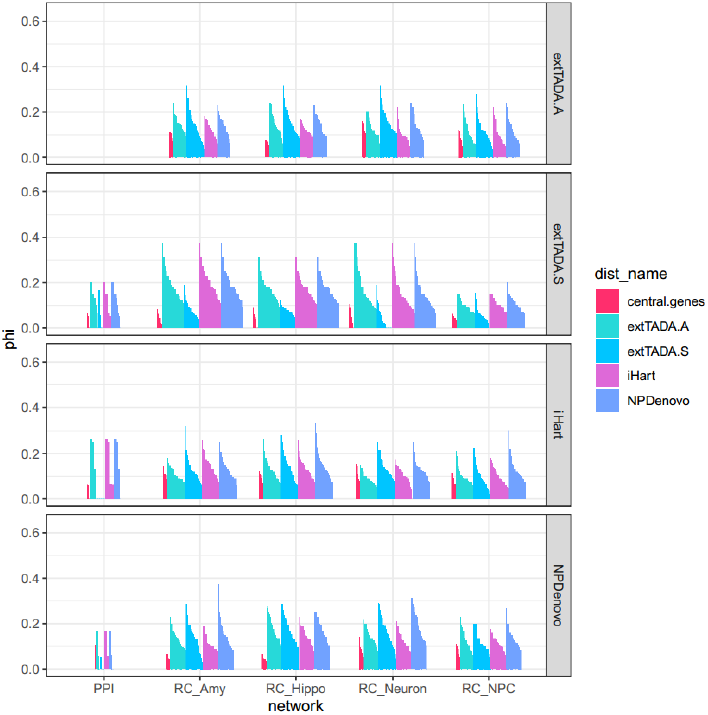
Plot of Phi statistics for InWeb brain PPI network (“PPI”) and four regulatorycircuits.org (“RC”) networks: Hippocampus (“Hippo”), amygdala (“Amy”), NEU+ neurons, astrocytes, and neuroprogenitor cells (“NPC”). Vertical breaks represent the study used to calculate phi, while the colors represent those studies used to define proposal core genes, or network central genes.

